# EyaHOST, a modular genetic system for investigation of intercellular and tumor-host interactions *in Drosophila melanogaster*

**DOI:** 10.1101/2024.09.06.611647

**Authors:** José Teles-Reis, Ashish Jain, Dan Liu, Rojyar Khezri, Sofia Micheli, Alicia Alfonso Gomez, Caroline Dillard, Tor Erik Rusten

## Abstract

Cell biology and genetic analysis of intracellular, intercellular and inter-organ interaction studies in animal models are key for understanding development, physiology, and disease. The MARCM technique can emulate tumor development by simultaneous clonal tumor suppressor loss-of-function generation coupled with GAL4-UAS-driven oncogene and marker expression, but the utility is limited for studying tumor-host interactions due to genetic constraints. To overcome this, we introduce EyaHOST, a novel system that replaces MARCM with the QF2-QUAS binary gene expression system under the *eya* promoter control, unleashing the fly community genome-wide GAL4-UAS driven tools to manipulate any host cells or tissue at scale. EyaHOST generates epithelial clones in the eye epithelium similar to MARCM. EyaHOST-driven Ras^V12^ oncogene overexpression coupled with scribble tumor suppressor knockdown recapitulates key cancer features, including systemic catabolic switching and organ wasting. We demonstrate effective tissue-specific manipulation of host compartments such as neighbouring epithelial cells, immune cells, fat body, and muscle using fly avatars with tissue-specific GAL4 drivers. Organ-specific inhibition of autophagy or stimulation of growth-signaling through PTEN knockdown in fat body or muscle prevents cachexia-like wasting. Additionally, we show that *Ras^V12^, scrib^RNAi^* tumors induce caspase-driven apoptosis in the epithelial microenvironment. Inhibition of apoptosis by p35 expression in the microenvironment promotes tumor growth. EyaHOST offers a versatile modular platform for dissecting tumor-host interactions and other mechanisms involving intercellular and inter-organ communication in *Drosophila*.

**Highlights:** \**eyes absent*, eye disc-specific enhancer drives clonal KD recombinase flip-out activated QF2 expression in the larval eye epithelium for simultaneous QUAS-driven gain and loss-of-function analysis of gene function.

*Clones are visualized by QUAS-tagBFP or QUAS-eGFP facilitating analysis of existing fluorescent reporters.

*The GAL4-UAS system and existing genome-wide genetic tools are released to independently manipulate any cell population in the animal for cell biology, intercellular or inter-organ analysis for developmental, physiological, or disease model analysis.

*Fly avatars for tumor-host interaction studies with multiple organs allow live monitoring and manipulation of tumors and organs in translucent larva.

## Introduction

The success of *Drosophila melanogaster* as a genetic model system to study development, physiology and disease hinges on the powerful techniques developed for mosaic clonal genetic gain and loss of function analysis paired with the fly community-generated ecosystem of genome-wide tools and stock collections ^1^. Mosaic loss-of-function genetic analysis was revolutionized by the introduction of precise reciprocal mitotic recombination between homologous chromosome arms, which can induce homozygosity of recessive alleles on the affected chromosome arm upon cell division. This is facilitated by expression of the yeast FLP recombinase that cut FLP-recombinase target (FRT) sites inserted close to chromosomal centromeric regions and promote chromosome arm swapping and repair (^2–4^ reviewed in ^5^). Directing FLP recombinase activity to the germline or the eye-antennal disc of the larvae further refined this technique and pioneered its use in organ-specific genetic analysis ^6–9^. The ability for directed tissue and time-controlled gain- and loss-of-function genetic analysis was achieved by introduction of the yeast GAL4 transcription factor, whereby any gene can be expressed under control of the GAL4 Upstream Activating Sequence (UAS)^10^. Genome-wide UAS-driven tools are now available from *Drosophila* stock collections to over-express, knock down, or knock out any gene using RNAi or CRISPR/Cas9 technology ^1^. Detecting FLP-FRT loss-of-function clones, which are typically small and situated within a vast pool of normal cells, was challenging and limited further manipulation possibilities. A solution to this was provided by the introduction of the Mosaic Analysis with a Repressible Cell Marker (MARCM) technique ^11,12^. In MARCM, effective positive labeling of loss of function clones generated by the FLP-FRT technique is achieved by combining the GAL4-UAS system and the FLP-FRT system. In MARCM, FLP serves two functions, activating GAL4 in a tissue of choice and mediating mitotic recombination. First, it excises a FRT-flanked stop cassette between the ubiquitously expressing *actin5C* promoter and GAL4 ensuring an inherited stable GAL4 expression in clonal descendants in the tissue where FLP is expressed providing tissue specificity. GAL4 activity is kept inactive by expression of the ubiquitously expressed GAL4 repressor, GAL80 inserted on the sister chromatid of choice for which mitotic recombination is to be performed. As a result, GAL80 is removed only in cells where mitotic recombination leads to daughter cells that have lost the GAL80 copy and simultaneously induced homozygosity of the sister chromatid arm carrying a recessive allele of the gene of interest. In sum, this allows positive labeling, visualized by UAS-GFP expression, of a clone for which loss of function of a given gene has occurred and can then be coupled with any GAL4-UAS driven manipulation of interest. In two seminal studies, researchers capitalized on this advantage to demonstrate that cells with simultaneous loss of a tumor suppressors and overexpression of an oncogene can emulate genetic cooperation and malignant tumor formation and progression of tumor-initiating cells in flies ^13,14^. Various tumor models ^15^ have been developed that display cancer hallmarks similar to human cancer ^16^ involving highly conserved signalling pathways ^17–19^. *Drosophila* is increasingly used as an *in vivo* system for studying complex interactions that venture beyond tumor-intrinsic mechanisms to probe reciprocal tumor-host interactions with the microenvironmental stromal compartment and systemic tissues during cancer progression and how this impacts tumor growth, metabolism, deteriorate host health and promotes death^20^.

One of the most studied *Drosophila* tumor models is a carcinoma model that arises from the cooperation between the oncogene *Ras^V12^* (K-RAS ortholog) and the loss of the tumor suppressor *scribble* (hScrib ortholog)^13,14^. The *Ras^V12^; scrib^-/-^* tumors are formed in a simple *Drosophila* larval epithelial organ, the eye-antennal imaginal disc (EAD), from randomly generated single cells using the MARCM technique under eyeless-FLP control (EyMARCM) ^8,13,14,21^. *Ras^V12^; scrib^-/-^* tumors rapidly develop to large tumors that display a complex set of malignant cancer hallmarks that mimics surprisingly complex aspects of human cancer biology, including systemic cancer cachexia-like paraneoplastic effects ^22–26^. *Ras^V12^; scrib^-/-^* tumors grow perpetually through cell cycle and cell growth stimulation and are resistant to apoptosis and differentiation. Tumor cells migrate from the EAD through extracellular matrix degradation and invade other neighboring larval organs. In this process, *Ras^V12^; scrib^-/-^* tumors also attract macrophage-like-cells hemocytes that accumulate at the tumor surface, and ultimately induce the wasting of energy storage organs such as the muscle and fat body. Studies have established that the autonomous programs controlling *Ras^V12^; scrib^-/-^* tumorigenesis involve a cocktail of active signalling pathways that are also known to be drivers of human carcinogenesis ^13,14,24,27–38^ (reviewed in ^18^).

More recently, studies have focused on tumor-host interactions using this carcinoma model and revealed mechanistic insights into how transformed cells engage with hemocytes, nearby epithelial cells or affect distant organs during tumor progression and execution of organ wasting ^24,28,37,39^. However, because the EyMARCM technique utilizes the GAL4-UAS system to form *Ras^V12^; scrib^-/-^* tumors, the way to manipulate other organs and cellular compartments during tumorigenesis mainly relied on classical LOF analysis and inheritance of mutant alleles in the non-tumor epithelial compartment generated upon mitotic recombination with the homologous *scrib mutant* chromosome ^28^ or fly-to-fly transplantation/transfusion techniques ^37,39^ from mutant animals. These approaches are laborious, preclude utilization of the genome-wide available tools of the GAL4-UAS system, and are further restricted by the physical location of a given allele analyzed and to the epithelial cell population of the eye antennal disc. Moreover, potential confounding effects due to transplantation, which creates a wound and potential infections, cannot be ruled out.

Alternative genetic systems of generating tumors independent of the GAL4-UAS system such as the lexA-lexAop system from *Escherichia Coli* ^40^ or the QF-QUAS system from *Neurospora crassa* ^41^ have recently been successfully adopted for inter-organ communication and paraneoplastic effect studies in response to tumor presence in adult and larval models ^23,25,42,43^. So far, these models are less restricted to a defined cell population and stop short of the dynamics and granularity provided by the clonal tumor-inducing clones of the EyMARCM system that we predict will impact early tumor development.

### Design

To overcome the technical limitations and caveats described above and create a model to study tumor-host interactions at scale, we set out to establish a tumor model in the *Drosophila* larva that ideally would have the following characteristics: 1) The tumor-generating system should be independent of the GAL4-UAS and FLP-FRT system to reserve these techniques for ready genetic manipulation of any host cellular compartment or organ at scale. 2) Tumor generation should be epithelial and clonal in origin to enable genetic analysis of interaction between tumor-initiating cells with neighboring epithelial cells, hemocytes and any other cells or organs. 3) Tumor generation needs to be stereotypic and quantifiable to allow assessment of effects on tumor size and progression, including tissue invasion as a result of manipulating tumor-host interaction. 4) Genetic tools need to be genetically “light” such that animals are healthy and sufficient chromosomes are available to load GAL4 drivers for specific cell populations and any tool for knockdown, overexpression or reporters can be easily incorporated by a single generation of breeding. This is distinct from the EyMARCM system where all autosomes are typically occupied. 5) The system should allow fluorescent labeling of tumor and host populations that are being genetically manipulated, allowing for direct observation and quantification without dissection when needed. 6) Tumor labeling by multiple fluorescent colours should be established to allow the ready use of available community-generated fluorescent reporters. 7) GAL80 must be implemented to enable time-controlled expression of GAL4-driven expression, and importantly, enable strict non-overlapping compartments of gene manipulation of tumor-initiating cells and neighboring epithelial cells for interclonal interaction studies. 8) Tumor generation should ideally be more restricted to the eye disc than the current EyMARCM system, which aside from the eye-antennal discs, also display tumor clone generation in male gonad primordia and cells of the optic lobes of the brain. 9) The system should be modular such that it can be readily adopted to model a diverse set of tumors by replacing tumor driver mutations, or to study other non-cancer related questions that can be modeled genetically.

We herein describe the combination of the QF2-QUAS binary expression system ^44^ to generate eye epithelium clonal *Ras^V12^; scrib^RNAi^*-driven tumor populations with the GAL4-UAS system utilized for host gene expression to create a highly modular genetic system for tumor-host studies which we term EyaHOST. We envision that the adoption of this system for tumor, inter-tissue and inter-organ communication will greatly facilitate tumor-host studies and other questions at scale.

## Results

### Design and validation of the components of the EyaHOST system

Although the EyMARCM system has been successfully used to study some aspects of tumor-host interactions, the utility is limited since all autosomes in the *Ras^V12^; scrib^-/-^* tumor model are utilized and the GAL4-UAS system is bound up for tumor generation and therefore not available to manipulate host compartments at scale (Fig1A). Other genetic manipulations, reporters and rescue constructs need to be custom made, recombined to existing chromosomes or located to the X chromosome.

**Figure 1.**
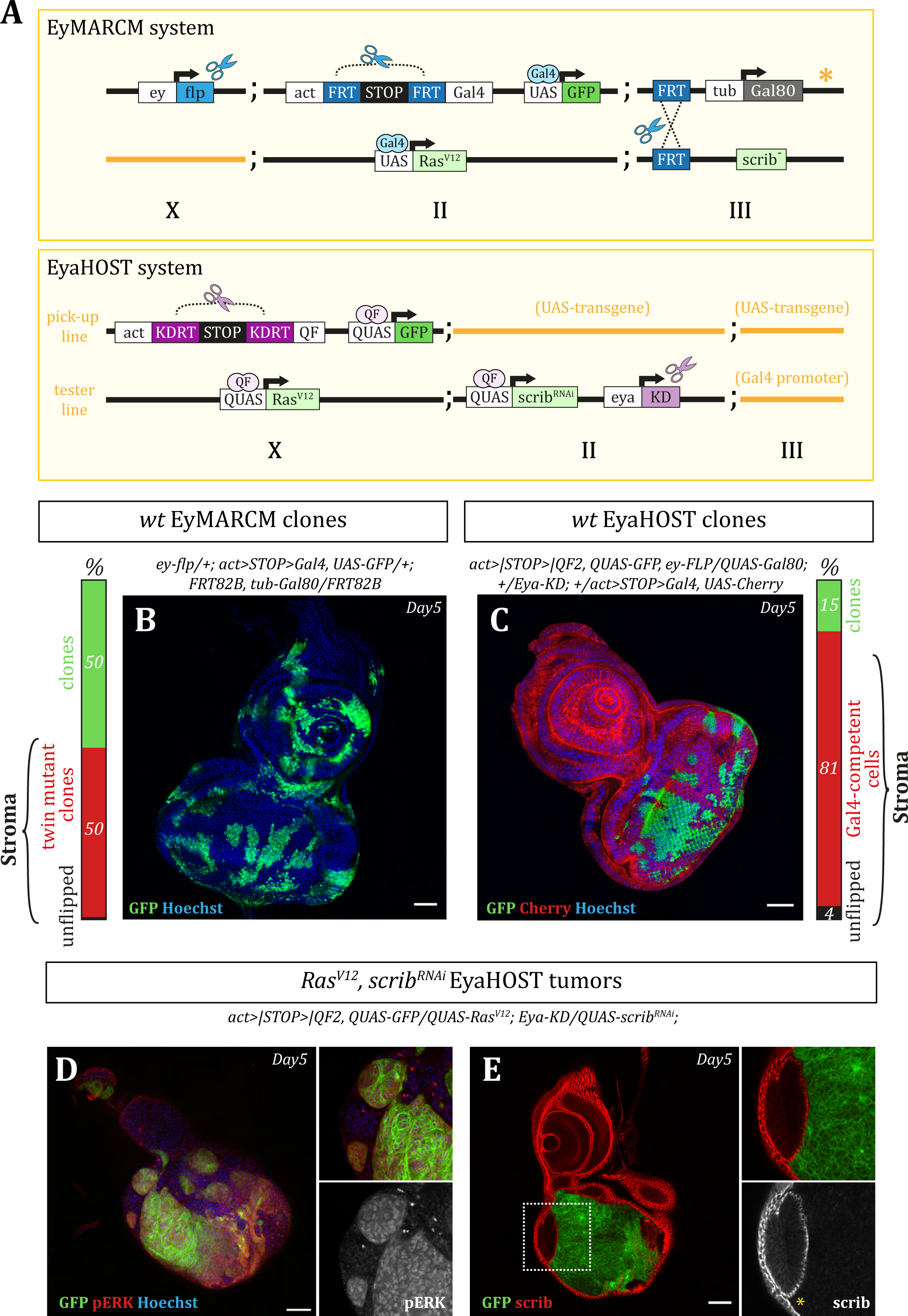
EyaHOST system allows the independent genetic manipulation of two cell populations in the EAD. (A) Diagram comparing the genetic design strategies of commonly used EAD clonal system EyMARCM and EyaHOST. The diagram shows the different transgenes inserted along the X, II, III chromosomes of *Drosophila*. Briefly, the EyMARCM system utilizes the FLP/FRT system and mitotic recombination for Ras^V12^ scrib^-/-^ tumour generation. Although highly effective, it requires four transgenes and occupies significant chromosome space, leaving only one homologous X chromosome available for additional genetic manipulations. In contrast, the EyaHOST system requires only two transgenes for clone generation, freeing one homologous II chromosome and both homologous III chromosomes for further genetic manipulation. Clone generation is achieved through tissue-specific activation of an Act-KDRT-STOP-QF cassette via the KD/KDRT system, driven by the Eya promoter. This system also employs the QF-QUAS system for tumor generation, which allows for host-independent manipulation using the widely adopted GAL4-UAS system. The EyaHOST system is activated by crossing the pickup and tester fly lines. The tester line is designed to have the III chromosome free for host tissue-specific GAL4 transgenes, while the pickup line can accommodate any desired RNAi, overexpression, reporter, or other transgenes on II and III chromosomes. (B) and (C) Confocal images and percentage of activated clonal wild-type cell populations in EAD of the EyMARCM and EyaHOST systems, respectively. Hoechst in shown in blue, GFP in green, Cherry in red. (C) As shown by Cherry expression, the EyaHOST system enables independent GAL4-UAS manipulation of neighboring cell populations. Specifically, an Act-FRT-STOP-FRT-GAL4 cassette is activated via the FLP/FRT system under the control of the Ey promoter, leading to GAL4 expression across the entire disc. Subsequently, QUAS-GAL80 expression inhibits GAL4 activity within the QF-expressing population. (D) GFP-positive Ras^V12^, scrib^RNAi^ tumours (green) generated through the EyaHOST system, stained for pERK (red), or E) stained for apical/basal polarity gene protein scrib (red). Samples shown are EAD of animals dissected at day 5 after egg-lay (AEL). Scale bar – 50 μm

To induce clonal expression and tumor generation in the eye antennal disc independent of GAL4-UAS and FRT system we chose the second generation QF2-QUAS binary gene expression system, optimized against cellular toxicity effects by QF, and the KD-KDRT recombinase system to generate inheritable gene expression ^41,45,46^. The KD recombinase was selected as it is non-toxic, and the consensus target site is furthest apart in length and sequence from the FRT target site among the tested and available site-specific recombinase systems established in flies ^45^. In order to free ourselves from the requirement of mitotic recombinations to generate homozygous *scrib^-/-^* cells, we turned to knock-down of *scrib* by creating a *20XQUAS-scrib^RNAi^* transgene, which efficiently decreases Scrib protein levels (Fig1E). We used an existing *5XQUAS-Ras^V12^* transgene (kind gift from P.Gallant) that constitutively activates dERK/MAPK signalling (Fig1D) and an available *5XQUAS-mcd8::GFP* to label clones and tumors (Fig 1D, E). To ensure strong and constant heritable expression of the QF2 transcription factor, we placed the QF2 gene under the control of the ubiquitous Act5C (act) promoter, previously used in the EyMARCM system (Fig1A). QF2 expression is maintained off by default, as we inserted an SV40 terminator STOP cassette between the act promoter and the QF2 open reading frame (ORF). The STOP cassette is flanked by two KD Recombinase Target sites (KDRT, further abbreviated >| which symbolizes an inverted K). To excise the *act>|STOP>|QF2* cassette and activate the Q-system in randomly selected EAD cells, the KD recombinase which recognizes KDRT sites has to be expressed and active only in a subset of EAD cells ^45^. The *eyeless* enhancer used in the previous EyMARCM system (Fig1A) is active in the entire EAD as well as in the optic lobes of the central nervous system and in male gonads (CNS) (^8^ and SuppFig1C). Therefore, we searched for another eye-specific enhancer that would have a more restricted expression pattern to direct expression of the KD recombinase. We selected an *eye absent* (*eya*) enhancer (kind gift from J.Kumar) and established *eya-KD* transgenes. Serendipitously, when used with the *act>|STOP>|QF2 stop cassette, eya-KD* triggers the formation of random clones with very similar dynamics to the EyMARCM without the need for mitotic recombinations used in the EyMARCM system (Fig1B,C). We named this new technique after the *eya* gene and the possibility to manipulate any cells of the host by another binary gene expression system, EyaHOST. EyaHOST clones always develop in the eye part of the EAD (100% EAD with clones in the eye part), often in the peripodium cell layer (74% EAD with peripodial clones) but rarely in the antennal part (23% EAD with antennal clones) whereas *wt* EyMARCM clones always form in every part of the EAD (Fig1B, C and SuppFig1). As a consequence, at the late larval stage, *wt* EyaHOST clones represent a smaller portion of the total EAD compared with *wt* EyMARCM clones (15% vs. 50% of the EAD respectively - Fig1B,C, SuppFig1B). However, similar clone occupancy is observed specifically in the eye region of the EAD for both the EyMARCM and EyaHOST systems (30% vs. 27.4% of eye region only - SuppFig1B). We never observed the formation of *wt* EyaHOST clones in the CNS (0%) compared with the EyMARCM system (100%), constituting a valuable improvement for studying migration of EAD tumor cells to the brain (SuppFig1C). Similarly, we never observed EyaHOST clones in male gonads (0%) while this is frequently observed using the EyMARCM system (100% - SuppFig1C). Activation of the EyMARCM system in the gonads has been a challenge for maintaining some stable EyMARCM *Drosophila* stocks. The *act>STOP>Gal4* (Fig1A) cassette occasionally lose the STOP cassette in the germline, and a constitutively expressed *act-Gal4* transgene results and is passed onto the next generation.

Because of the potential for stop cassette loss, we decided as a general principle, to avoid placing a recombinase and its corresponding *act>|STOP>|QF2* cassette in the same *Drosophila* line. Therefore, the *eya-KD* transgene has been placed in the “tester line” and the *act>|STOP>|QF2* cassette in the “pick-up” line (Fig1A). Small EyaHOST clones also form in the larva mouth hook (100%) whereas we observed only few clones in this location using the EyMARCM system (16%). We also observed sporadic small EyaHOST clones in the larva body wall but not in the EyMARCM system (SuppFig1C). At lower frequencies, clones form in the trachea, fat body, and salivary glands but never in the other imaginal discs was found using either system, with the exception of genital discs which occasionally form clones with the EyMARCM system (SuppFig1C). We found that EyaHOST clone generation of the can be affected by the temperature at which the larva are reared. We observed a 2.2 fold increase in clone volume when larvae developed at 18C compared to 25C, and a 4.5 fold decrease in clone volume when raised at 29C compared to 25C (SuppFig3A). This suggests that the KD-KDRT system may be temperature sensitive, which to the best of our knowledge has not been demonstrated.

Overall, the EyaHOST system reliably and stereotypically generates clones in the eye part of the eye-antennal disc with a clone density distribution and coverage similar to the EyMARCM system, whereas the total clone density and coverage in the whole EAD is 4-fold smaller. Sporadic clone generation is active in fewer other organs compared to the EyMARCM system (SuppFig1C) and never in the germ line, genital disc and CNS.

### *Ras^V12^; scrib^RNAi^* EyaHOST tumors display the hallmarks of the *Ras^V12^, scrib^-/-^* EyMARCM tumors

After validating the different transgenes required for clonal induction, we went on to assess whether we could generate and recapitulate the properties of *Ras^V12^; scrib^-/-^* EyMARCM tumors utilizing the EyaHOST and the Q-system transgenes *5XQUAS-Ras^V12^ and 20xQUAS-scrib^RNAi^*. The *Ras^V12^; scrib^-/-^* hallmarks are described in ^18^ and include: massive tumor growth, developmental delay, differentiation inhibition, migration, muscle wasting, fat body wasting and non-autonomous autophagy induction.

EyaHOST tumors grow large and ultimately take over the whole anterior part of the larva similarly to EyMARCM tumors (SuppFig2A, Fig2A). However, EyaHOST tumors reach the size of Day 10 after egg-lay (AEL) EyMARCM tumors after 18 Days AEL (Fig2A). This paucity stems from the fact that the pool of EAD cells from which the EyaHOST tumors originate is smaller than in the EyMARCM system (Fig1B,C).

**Figure 2.**
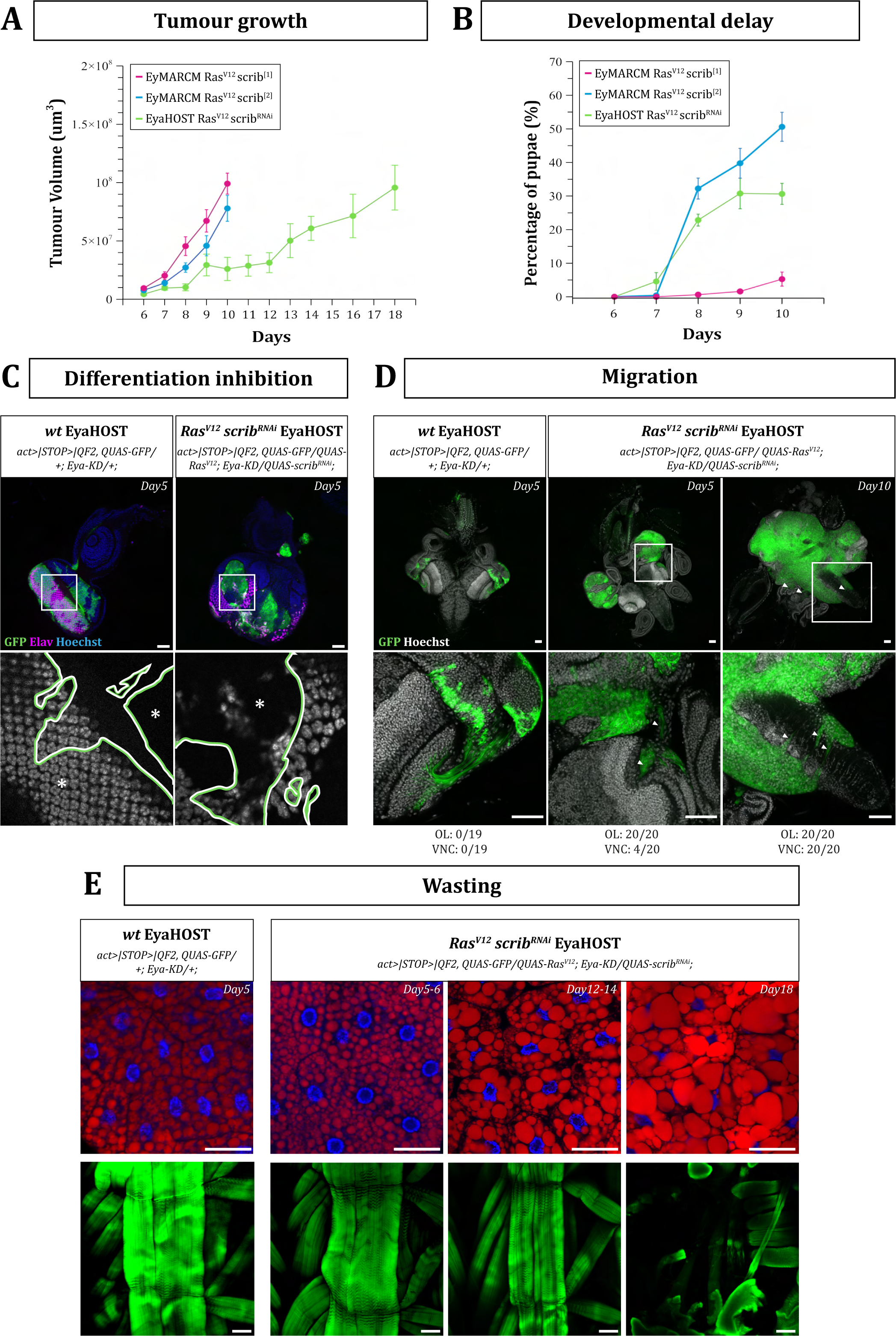
EyaHOST Ras^V12^, scrib^RNAi^ tumours recapitulate known tumour hallmarks and paraneoplastic effects. (A) Ras^V12^, scrib tumour volume changes over time for the EyMARCM system using two different scrib alleles, scrib^1^ (pink), scrib^2^ (blue), as well as for EyaHOST tumours (green). (B) Pupariation percentage comparison between EyMARCM system, scrib^1^ (pink), scrib^2^ (blue) alleles, and EyaHOST animals (green). (C) Confocal images of EAD with GFP positive WT clones (left and green) or Ras^V12^, scrib^RNAi^ tumours (right and green), stained for Hoechst (blue) and Elav (pink). Scale bar - 50 μm In the close-up image, the clone border is marked by a line, with green indicating the clone side and white representing the neighbouring cells. Asterisk marks the inside of the clone. (D) Microscope images of larval cephalic complexes (Mouth hook, EAD, CNS) from WT at D5 AEL and Ras^V12^, scrib^RNAi^ tumours at D5 or 10 AEL. Quantification shows the frequency of invasion of GFP-positive cells in the optic lobe (OL) and ventral nerve cord (VNC). Scale bar - 50 μm Insets demonstrate the absence of GFP-positive cell nuclei in the OL at D5 in WT, contrasted with their presence in Ras^V12^, scrib^RNAi^ tumours at D5. By D10, the optic lobes are completely invaded by Ras^V12^, scrib^RNAi^ tumour cells, which also extend into the VNC. (E) Confocal images of the fat body (top) and muscle (bottom) of WT D5, and Ras^V12^, scrib^RNAi^ tumours at D5, D12-14, and D18 animals. Fat body preparations were stained for LipidTOX (red) and Hoechst (blue), and muscle fillets were stained for actin using phalloidin 594 conjugate. Scale bar - 50 μm

*Ras^V12^; scrib^-/-^* EyMARCM tumor-bearing larvae suffer from delayed entry to pupal stage and most will linger as large (giant) larvae that eventually die. The remaining larvae are able to pupariate but die during metamorphosis before reaching adulthood. Therefore, *Ras^V12^; scrib^-/-^* EyMARCM tumor bearing larvae have a low pupariation rate. We compared the pupariation rates of the *Ras^V12^, scrib^RNAi^* EyaHOST tumor-bearing larvae with the pupariation rate of *Ras^V12^; scrib^-/-^* tumor bearing larvae using two different versions of the EyMARCM system: “EyMARCM“ (the most widely used system) and “EyMARCM^2^”. These two EyMARCM systems employ different *UAS-Ras^V12^* transgenes and different *scrib* loss-of-function alleles (*scrib^1^* and *scrib^2^* resepectively). At Day 10 AEL, the EyMARCM pupariation rate is very low (around 5%) and the EyMARCM^2^ pupariation rate is much higher (around 50%) while the EyaHOST pupariation rate is around 30% (Fig2B). Thus, EyaHOST tumors trigger a similar delay of pupation to that of EyMARCM and EyMARCM^2^ tumors. We observed that the pupariation rate of EyaHOST tumor bearing larvae is affected by the population density unlike EyMARCM tumor bearing larvae. As a consequence, growing conditions of EyaHOST tumor bearing larvae have to be optimized in order to ensure the abundance of late staged lingering tumor-bearing larvae.

Similar to *Ras^V12^, scrib^-/-^* EyMARCM tumors, neuronal differentiation is repressed in *Ras^V12^, scrib^RNAi^* EyaHOST tumor, where the pan-neuronal marker Elav is seldom observed compared with *wt* EyaHOST clones (Fig2C). *Ras^V12^, scrib^RNAi^* EyaHOST tumors displayed cells with a stretched cell morphology, suggesting that they represent migrating cells. From day 5 AEL, tumor cells leaving the EAD and migrating along the optic nerve toward the optic lobes of the CNS are observed. At day 10, cells invade the surface and the inside of the ventral nerve cord in the CNS (Fig2D).

Systemically, at day 8 AEL, EyaHOST tumor-bearing larvae display typical fat body lipid droplet remodeling and morphological changes previously described as fat body wasting (Fig2E, Fig4D). They later undergo muscle wasting (Fig2E, Fig4C) characterized by muscle fiber shrinkage ^23,24,47^.

As previously described with the EyMARCM system ^24,28^, the growth of *Ras^V12^, scrib^RNAi^* EyaHOST tumors also induce an autophagic response in the surrounding EAD tissue, muscles, fat body and gut as shown by the accumulation of Cherry::Atg8a puncta in these tissues (Fig5A).

We next verified whether the previously described signalling pathways that are ectopically induced in *Ras^V12^, scrib^RNAi^* tumors were active in the EyaHOST tumors. We could clearly observe JNK, Jak/Stat and Yki signalling activation assessed in Day 5 AEL EyaHOST tumors with pJNK immunostainings (Fig3A), the *10XSTAT-GFP* (Fig3B) and the Kibra-LacZ (Fig3C) reporters, respectively.

**Figure 3.**
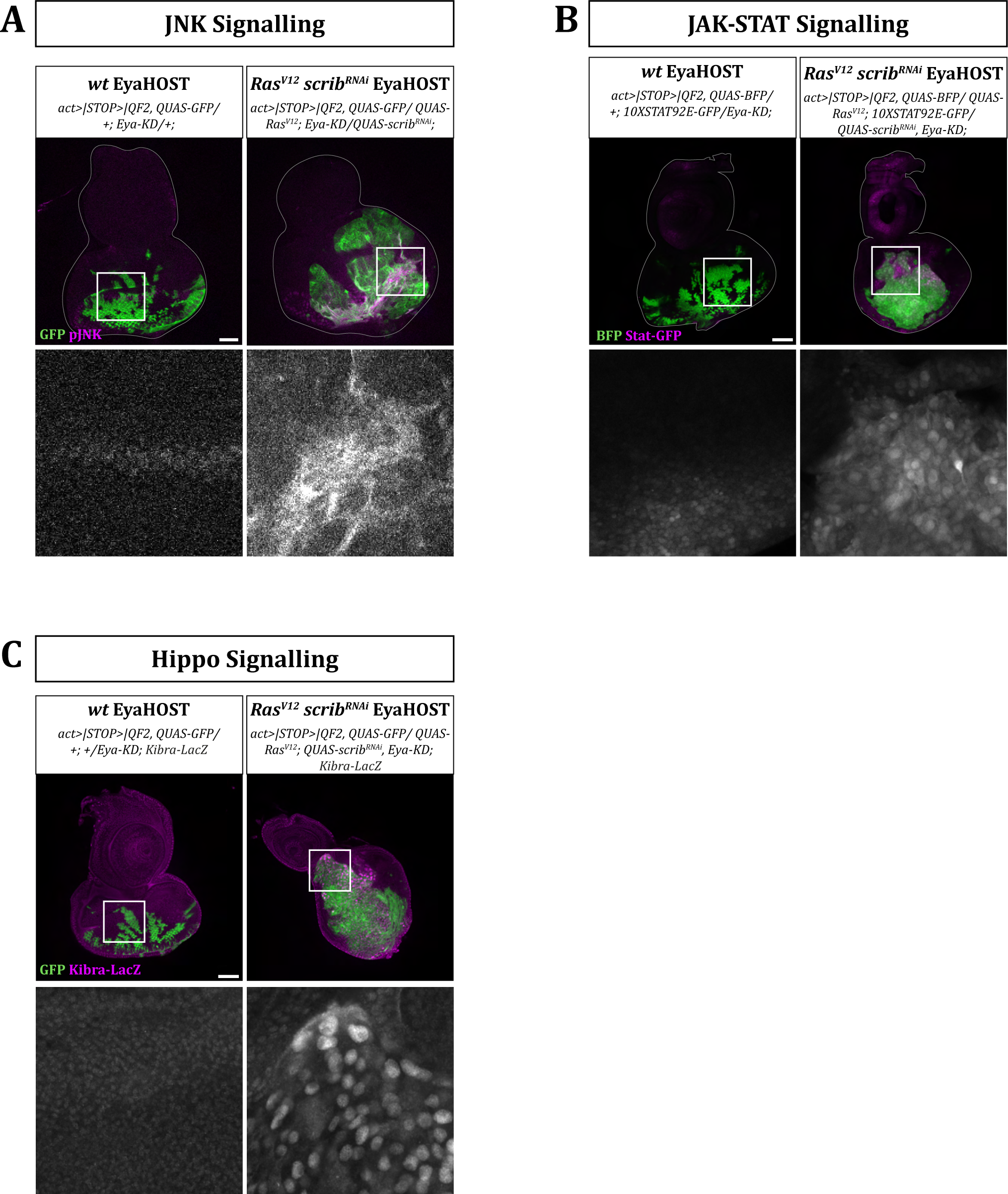
EyaHOST Ras^V12^, scrib^RNAi^ tumours recapitulate the known ectopic activation of the JNK, JAK-STAT, and Hippo pathway. In each set of panels from (A-C) shows confocal images of EAD from WT and Ras^V12^, scrib^RNAi^ EyaHOST larvae at D5. GFP (green) marks WT cells or Ras^V12^, scrib^RNAi^ EyaHOST tumour cells, while pathway-specific markers pJNK, Stat-GFP, Kibr-LacZ (pink) highlight the activation of the JNK, JAK-STAT, and Hippo pathways, respectively. Scale bar - 50 μm.

To conclude, despite the initial tumor cell population being smaller in the EyaHOST system, all known previously described phenotypes and cell signaling pathway changes unfold faithfully, opening up for mechanistic studies of tumor-host interactions.

### A modular tool to investigate tumor-host interactions

The EyaHOST system is genetically lighter given that three chromosomes are free to carry additional transgenes versus a unique X chromosome within the EyMARCM technique (Fig1A). We took advantage of this free space to build different EyaHOST fly avatars (Fig 4) that allow GAL4-driven genetic manipulation of different organs. To demonstrate the usability for manipulating these tissues in a tumor context, we forced the expression of UAS-Cherry in various organs of these EyaHOST avatars.

**Figure 4.**
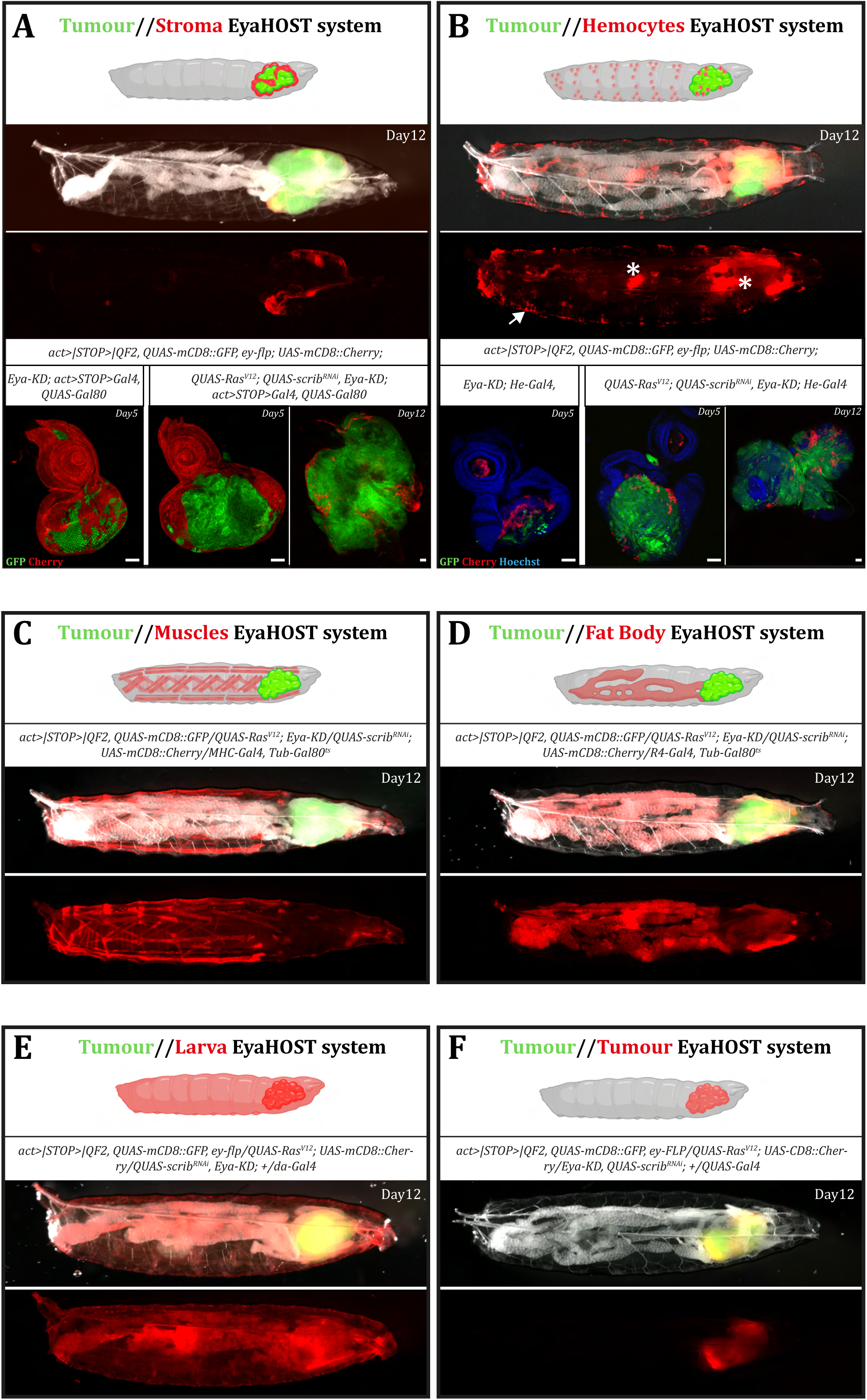
The modularity of the EyaHOST system enables the independent manipulation of any host tissue within a tumour context. Pannels (A-F) show stereoscope images of differently built EyaHOST avatars with QF-QUAS driven Ras^V12^, scrib^RNAi^ tumour bearing larva at D12. These avatars utilize GAL4-UAS genetic manipulation to target specific host tissues, as indicated by Cherry expression (red): (A) neighboring stromal cells, (B) hemocytes, (C) muscle, (D) fat body, and (E) the entire organism. Panel (F) demonstrates a tool designed for autonomous tumour manipulation within the same genetic background. Insets in panels (A) and (B) display WT D5 EAD, and D5 and D12 tumours from the stromal and hemocyte avatars, respectively. Asterisk in pannel (B) indicate ectopic expression of the Hemese driver in the gut and salivary glands. Arrow points to hemocyte Scale bar - 50 μm.

First, we created a tool that allows the genetic manipulation of the tumor-surrounding healthy EAD cells (hereafter referred as stroma). To do so, we repurposed some of the EyMARCM components for GAL4-stroma expression. Indeed, as the *eyeless* enhancer used in the EyMARCM system is active in the entire EAD, the FLP recombinase under its control can excise the *act>STOP>Gal4* cassette in the entire EAD (Fig1A). We thus used these two transgenes, *ey-flp* and *act>STOP>Gal4,* as the core components of the “**EyaHOST/Gal4-stroma**” avatar. Finally, in order to avoid GAL4 activity in the QF2-driven tumors and to allow dual compartment independent manipulation, we triggered expression of the GAL4 repressor GAL80 specifically in the tumors by adding a *20XQUAS-GAL80* transgene. Using *UAS-Cherry* expression as a proxy for GAL4 activity, we obtained strictly non-overlapping GFP^+^ Cherry^-^ tumor/ GFP^-^ Cherry^+^ stroma compartments (Fig 4A). To demonstrate we can specifically knockdown genes of interest exclusively in the surrounding GAL4 stroma, we downregulated the *white* gene, responsible for eye pigmentation, only in the neighboring cells. Indeed, we obtained adult *Drosophila* eyes that showed patches of pigmentation that coincided exclusively with the GAL4 Cherry^+^ compartment (SuppFig4A).

*Drosophila* macrophage-like cells, called hemocytes, are known to attach at the surface of *Ras^V12^, scrib^-/-^* EyMARCM tumors, but it has been challenging to investigate the molecular dialog between tumors and hemocytes in this context, without resorting to complex and less physiological techniques such as hemocytes transfusion and tumor transplantations ^37,39,48^. To facilitate, tumor-hemocyte interaction investigation, we built an “**EyaHOST/Gal4 hemocyte**” avatar. To do so, we added to the EyaHOST system the pan-hemocyte Gal4 driver, *He-Gal4,* and could observe Cherry^+^ hemocytes along the body wall of tumor-bearing larva (Fig4B, arrow), Cherry^+^ tumor-associated hemocytes at early and late tumor stages (Fig4B) as well as Cherry^+^ EAD resident hemocytes of *wt* EyaHOST clone-bearing larvae (Fig4B). It is important to note that strong Cherry levels is also seen in the salivary glands and parts of the gut (Fig4B, asterisks).

Previous studies in our lab and others described the progressive muscle and fat body wasting during *Ras^V12^; scrib^-/-^* tumorigenesis ^23,24,47^. To investigate this process, we generated an “**EyaHOST/Gal4 Muscle**” avatar that allows GAL4 expression in the muscle in a time-dependent manner. To do so, we used the muscle driver *MHC-Gal4* in combination with the ubiquitously expressed temperature sensitive Gal80 repressor, *tub-gal80^ts^*. At permissive temperature, we could clearly see wasting Cherry^+^ muscles in Day 12 AEL EyaHOST tumor-bearing larvae (Fig4C, SuppFig 4).

Following the same principle, we built an “**EyaHOST/Gal4 Fat Body**” avatar for time controlled genetic manipulation of the fat body in EyaHOST tumor-bearing larvae. We recombined the fat body GAL4 driver, *R4-gal4* with *tub-gal80^ts^* and could label the adipose tissue with Cherry at permissive temperature (Fig4D). In this setting, we also observed Cherry in the salivary glands (Fig4D, arrow).

Next, we designed a tool that renders possible GAL4-driven manipulation of the entire larva. Our first attempt using the ubiquitous *act-gal4* driver failed, likely because of toxic effects from expressing of GAL4 at very high levels throughout the entire organism. We then turned to milder ubiquitous driver and eventually used the *daughterless-gal4* driver for our “**EyaHOST/Gal4 larva**” avatar (Fig4E).

Finally, in investigating the communication axis between the tumor and a specific organ, it will be necessary to genetically manipulate the tumor itself within the same genetic background. A limited list of QUAS-transgenes is available so far. Although, it might expand dramatically in the coming years ^49^, we decided to use the Gal4/UAS system in order to also manipulate the tumor in our EyaHOST system. Therefore, we also created an “**EyaHOST/GAL4-tumor**” avatar, by incorporating an available *5XQUAS-Gal4* transgene (BDSC), which enables the expression of GAL4 only in EyaHOST tumors. This way, it is possible to drive the expression of any UAS-transgene in the EyaHOST tumors (Fig4F).

Overall, we have developed multiple EyaHOST avatar versions that allow the specific interrogation of the molecular crosstalk between *Ras^V12^, scrib^RNAi^* tumors and the stroma, hemocytes, muscle, fat body and whole larva. To enhance the system’s modularity and facilitate the easy swapping of GAL4 drivers, not only in the *Ras^V12^, scrib^RNAi^* EyaHOST system, but also in its corresponding controls: *wt* EyaHOST (Fig1C), *Ras^V12^* EyaHOST and *scrib^RNAi^* EyaHOST, we created in-house control versions as well as balanced versions for the fast loading of any driver/transgene (see table1 for all lines available).

### Host specific genetic manipulation with the EyaHOST system prevents tissue atrophy

We have previously shown that *Ras^V12^; scrib^-/-^* EyMARCM tumors induce autophagy activation in the stroma, muscle, fat body, gut, and that both local and distant autophagy activation contribute to tumor development and cachexia ^24,28^. The demonstration of the role of autophagy in these different tissues using the EyMARCM system has been genetically challenging. Indeed, it required to introduce *atg13^-^* or *atg14^-^* alleles to the EyMARCM system on the opposite *scrib^-^* homologous chromosome (Fig1B, asterix). As reported here, EyaHOST tumors trigger similarly autophagy in the stroma, muscle, fat body and gut (Fig5A). To showcase the capability of our various EyaHOST versions to easily investigate tumor-host interactions, a process that has been so far technically arduous to study, we inhibited autophagy by knocking down *atg1* in the fat body and muscle. In both scenarios, we observed the predicted rescue from tissue atrophy (Fig5B,C). Cancer cachexia is known to be caused in part by a metabolic shift from anabolism to catabolism of peripheral tissues caused by a depression of insulin signaling through expression of the Insulin Growth Factor Binding Protein (IGFBP) homolog *ImpL2* ^26,50^. To show we could invert this metabolic shift in peripheral tissues, we downregulated *pten* to stimulate the insulin signaling pathway. Indeed, we similarly observed a rescue from tissue atrophy (Fig5B,C, SuppFig5) as observed upon autophagy inhibition.

**Figure 5.**
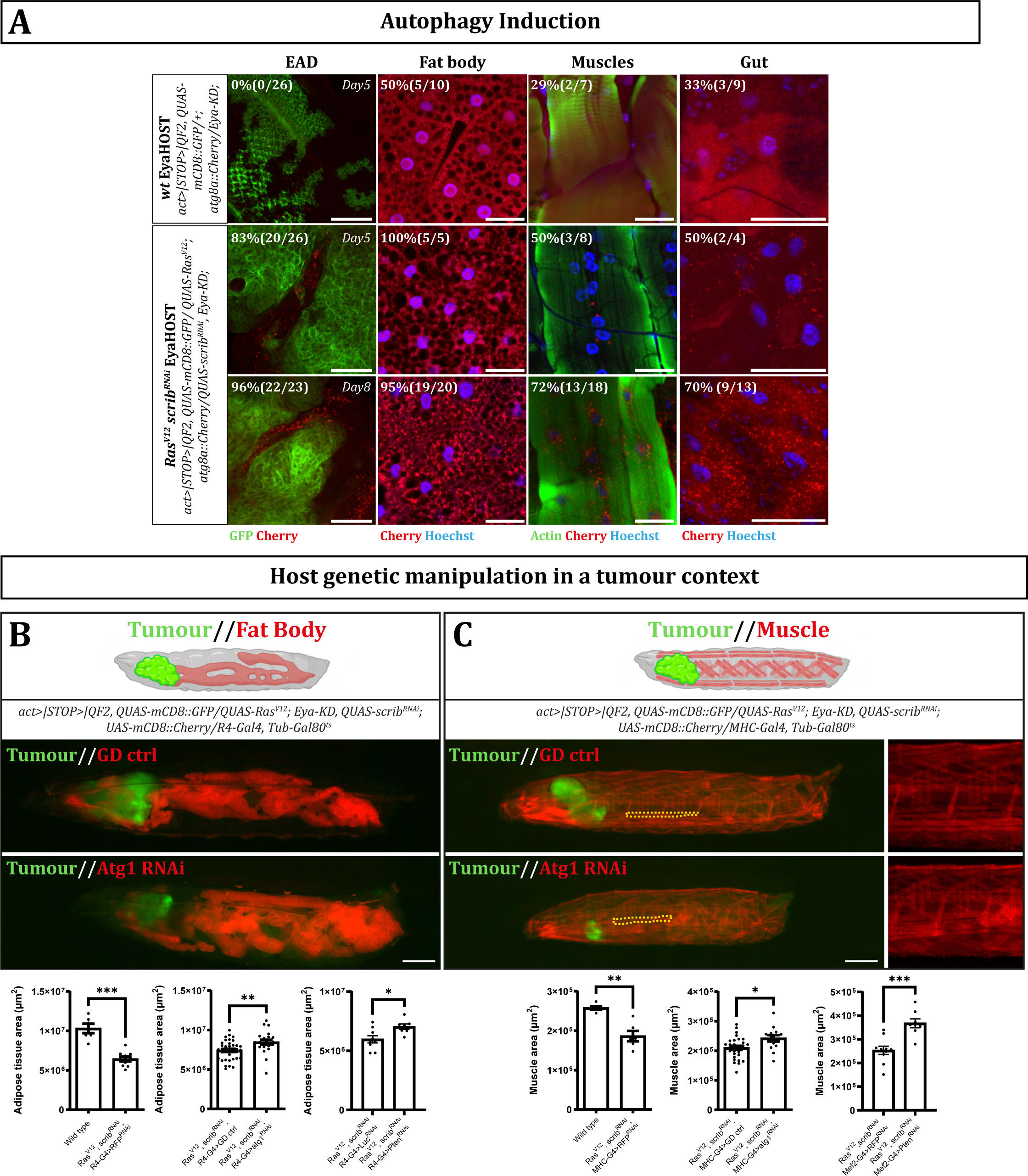
Tumour-induced host autophagy is recapitulated in the EyaHOST system and organ-specific autophagy inhibition prevents tissue atrophy. (A) Representative confocal images of EAD, fat body, muscles, and Gut in WT D5 larvae and Ras^V12^, scrib^RNAi^ tumour-bearing larvae at D5 and D8. Autophagy induction is indicated by ChAtg8 puncta (red) in various tissues. GFP-labelled Ras^V12^, scrib^RNAi^ tumours (green) are shown in the EAD column. Samples are counterstained with Hoechst (blue) for nuclei and actin (green) for muscle tissue. The percentage of samples showing ChAtg8 puncta is quantified for each tissue type. Scale bar - 50 μm (B) and (C) stereoscope images of tumour-bearing larvae with Atg1 knockdown in the fat body (B) and muscle (C), respectively. Quantifications of adipose tissue area and muscle area are provided below the images. Adipose tissue area and muscle area quantifications are also shown for pten knockdown in fat body and muscle (images shown in supplementary. fig 5). Insets in (C) show a close up of muscle structures. Statistical significance between groups was determined using the Mann-Whitney U test: *p ≤ 0.0332, **p ≤ 0.0021, ***p ≤ 0.0002. Scale bar - 700 μm.

In summary, as proof-of-concept, we demonstrate that the EyaHOST tool enables precise genetic manipulation of peripheral tissues within a tumor context, allowing us to experimentally address phenotypes relevant for tumor-host interaction studies.

### Microenvironmental undead cells enhances tumor growth

Previous studies have demonstrated that tumors can induce the death of neighboring healthy epithelial cells, paralleling interface behaviors observed during cell competition ^51–54^. This prompted us to investigate whether we could recapitulate and study this tumor-host interaction phenotype using the EyaHOST tool. Indeed, upon *Ras^V12^, scrib^RNAi^* tumor induction, we observed an increase in the microenvironmental programmed cell death detected by cleaved Death caspase-1 (Dcp1) immunolabeling (Fig 6A). Apoptosis, a highly regulated process, involves the coordinated action of three major classes of genes: pro-apoptotic genes (hid, rpr, grim), initiator caspases (Dronc and Dredd), and effector caspases (Drice and Dcp1)^55,56^. Several studies have shown that animal tissues have a regenerative capacity in response to apoptosis, where a dying cell will initiate a senescence-like program and secrete mitogens to promote its replacement through the proliferation of the neighboring cells. This process is termed apoptosis-induced proliferation (AiP) and extensive contribution to the discovery and study of this phenomenon has come from *Drosophila* studies ^57^. The great majority of studies addressing the mechanisms of AiP use a model where cell death is enforced genetically through overexpression of hid, but simultaneously effector caspases drice and dcp1 are blocked through the baculovirus antiapoptotic p35 protein. This generates “undead” cells that still have functional Dronc initiator caspase activity, which has been shown to be responsible for the inflammatory AiP phenotype ^57^. Interestingly, when using the EyaHOST model, we observed that when cell death was blocked through p35 in the neighboring cells of the WT population, the QF2 WT cell population expanded (Fig6A). Unlike shown in previous models of AiP where phenotypes are observed only upon forcing cell death genetically, we observed that the generation of undead cells from the cells targeted for developmental death was sufficient to observe a non-autonomous effect on clone expansion. Furthermore, we observed that the generation of undead cells in the microenvironment of *Ras^V12^, scrib^RNAi^* tumors promoted tumor growth (Fig6A). At the moment we have not addressed which are the communicating factors triggering this cooperativity. Furthermore, we were able to observe the undead cells in the microenvironment through cDcp1 staining. These contained high cDcp1 cytoplasmic levels, yet did not undergo pyknosis and apoptotic body formation. This observation matches the molecular activity of p35, which only blocks effector caspase activity after their processing by initiator caspases (Fig6A, yellow arrows) ^58^, and thus cDcp1 should remain accumulated in the cytoplasm of undead cells.

**Figure 6.**
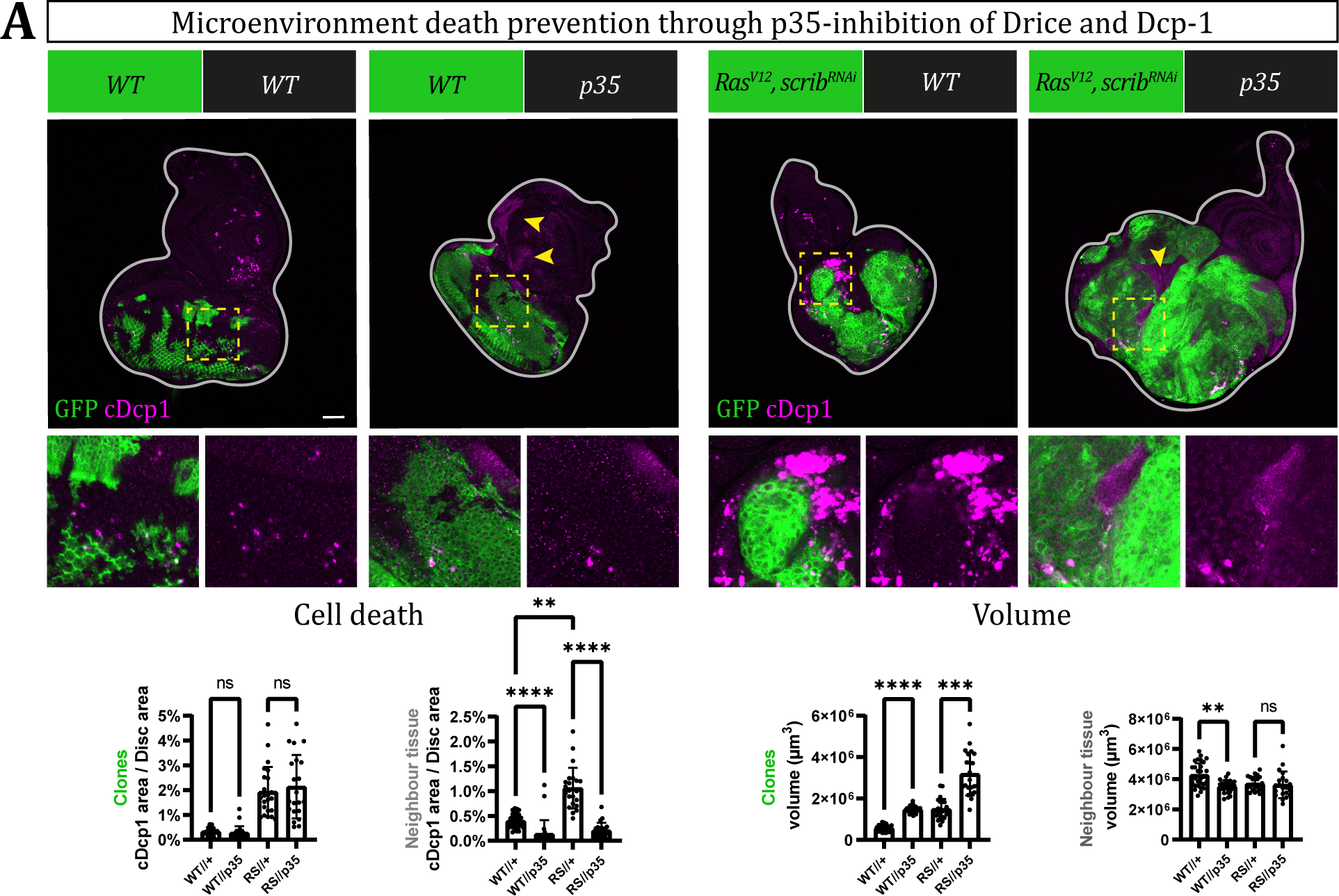
Microenvironmental generation of p35-mediated undead cells promotes tumour development. (A) Microenvironmental death prevention through p35-inhibition of Drice and Dcp-1. Representative microscope images of EAD at D5 of control and p35 microenvironment overexpression for WT and Ras^V12^, scrib^RNAi^ tumour conditions. WT clones and tumours are GFP-labelled (green) and stained for cDcp1 (magenta). Insets provide magnified views of the regions marked with dashed lines. Quantification of cell death, shown as the percentage of cDcp1-positive EAD area in clones and neighbouring tissue, is presented alongside volume measurements of tumour clones and adjacent tissue. Statistical significant differences between groups was assessed with a Kruskal-Wallis and post-hoc Dunn’s multiple comparison test: not significant (ns), *p ≤ 0.0332, **p ≤ 0.0021, ***p ≤ 0.0002, ****p ≤ 0.000. Scale bar - 50 μm.

Overall, these findings highlight the strength of the EyaHOST system in revealing the non-autonomous effects of cell death on tumor growth and offer significant potential for exploring the mechanisms of intercompartmental communication in both wildtype and tumor contexts.

## Discussion

### A new tool for systematic tumor-host interactions study in *Drosophila melanogaster*

Recent development of alternative binary expression systems (LexA-LexAop and QF-QUAS) and recombinase systems (FLP/FRT, KD/KDRT, RSR/RSRT, B2/B2RT, B3/B2RT) in *Drosophila melanogaster* unlocks the engineering of innovative and effective combinatorial uses, pushing back the technical limitations encountered in many fields of *Drosophila* research ^41,45,49,59^. We took advantage of these advances to generate EyaHOST, that allows the high-throughput and efficient investigation of the molecular crosstalks orchestrating tumor-host interactions or other biological questions. By repurposing the QF2-QUAS and the KD-KDRT recombinase systems for generating *Ras^V12^, scrib^RNAi^* tumors in the EAD specifically, we freed up the conventional genome wide GAL4-UAS and FLP/FRT systems for genetic manipulation of any organ or cell compartment of interest. We designed EyaHOST to be highly modular by ensuring minimal genetic occupancy on autosomes, facilitating the loading of any transgene of interest. We developed multiple fly avatars for manipulating the epithelial stroma, hemocytes, muscle, fat body, whole-larva, and tumor. Additionally, we generated the corresponding control lines (WT, scrib^RNAi^ only, and Ras^V12^), along with balanced lines, to enable the easy and rapid development of other combinatorial avatars. The EyaHOST system develop similar hallmarks to conventional EyMARCM tumors and activate the same key signalling pathways known to drive tumorigenesis in this context. As proofs-of-concept, we inhibited autophagy systemically in either muscle or fat body and observed a predicted rescue of observed organ wasting previously suggested by whole body mutant backgrounds. Furthermore, locally, we specifically blocked cell death in the stroma, which resulted in enhanced tumor growth. This observation aligns with previous findings on apoptosis-induced proliferation (AiP). However, our study reveals that the generation of undead cells, either from those destined for normal developmental cell death or from cells culled in the microenvironment due to the presence of the tumor, is sufficient to drive non-autonomous tumor growth.

Other genetic systems have been developed to create tumors in the EAD and manipulate other organs or compartments which were elegant but restricted to one type of tumor-host interactions. For instance, the QF/FLP genetic system developed by Lodge *et al.* in 2021 ^23^ enabled the generation of *QUAS-Ras^V12^, QUAS-scrib^RNAi^* tumors in the entire EAD using the conventional EyMARCM-derived *Ey-flp* transgene excising a tailored *act>STOP>QF2* cassette. This system was combined with the *MHC-Gal4* and *R4-Gal4* drivers for the genetic manipulation of the muscle and fat body, respectively. It elegantly allowed the study and identification of key factors involved in systemic organ wasting. In contrast to EyaHOST, tumors are in this case generated in the complete EAD and optical lobe, foregoing detailed studies of tumor/microenvironment interactions with cell populations such as neurons, glia, hemocytes and epithelial compartments. For these reasons, this system is optimal for studying systemic interactions in a candidate-based approach.

Enomoto *et al.* utilized another genetic system based on the combination of the QF binary expression system and the EyMARCM system to study local interactions between QF-driven *Ras^V12^* tumors and *Src*-overexpressing EyMARCM clones in the EAD ^60^. This technique allowed the discovery of the oncogenic interclonal cooperation between *Ras^V12^* and the oncogene *Src* to form malignant tumors.

Because it uses the EyMARCM technique, this system presents the disadvantages cited previously: a heavy genetic burden on all autosomes, limiting the execution of big scale genetic screenings and the unwanted expression in the gonads.

We believe that the EyaHOST system represents a great addition to these existing elegant techniques as it provides a high modularity for systemic and local studies of tumor/host interactions, genetic stability and healthiness of the avatar fly lines, and the possibility to execute large scale screening experiments of different organs with the same genetic background and appropriate controls.

### Limitations of the study

The robustness of the results obtained working with the EyaHOST system depends greatly on the strength, stability and specificity of the GAL4 driver used to genetically manipulate an organ of choice. For instance, the Tumor-hemocyte EyaHOST avatar effectively allows the genetic manipulation of larval hemocytes. However, the *He-Gal4* driver is also active in the salivary glands and parts of the gut. Therefore, any phenotype observed upon genetic manipulation of the hemocyte using this version of the EyaHOST system will need to be carefully investigated to distinguish the contribution of the hemocytes from the contribution of the salivary gland or guts to the phenotype observed. Carefully picking a tight organ-specific Gal4 driver or controlling for unwanted expression pattern would be crucial to avoid bias in the studies. Improving the expression pattern of GAL4 drivers have been a long time concern in the fly community and techniques such as the split-GAL4 system might be used to optimise intersections and refine Gal4-controlled expression patterns ^61^.

As stated previously, the percentage of clones activated in the EAD of the EyaHOST tumor-bearing larvae seems to impact the pupariation rate. We observed that a high larval crowdedness correlates with a high number of lingering larvae. Therefore, to ensure sufficient number of larvae for analysis. past Day5 AEL, culture conditions should be optimized in each laboratory condition without provoking starvation-induced developmental delay. We have optimized these growing conditions empirically for most EyaHOST avatars and generally perform biological replicates in parallels to limit variation.

Finally, although expanding the range of tumor/host interactions studies to both local and systemic interactions, the EyaHost technique is for now strictly limited to the EAD *Ras^V12^ scrib^RNAi^* (*Ras^V12^* only, *scrib^RNAi^* only and *wt*) tumor model. In future work it can be easily expanded to other genetically tailored tumor types and/or primary locations. Tumor type could be modified by replacing the *QUAS-Ras^V12^* and *QUAS-scrib^RNAi^* transgenes by any other relevant QUAS-transgene, whose number keeps increasing over time thanks to the work of several labs ^41,44,49,62^. Tumor location is driven by the *Eya-KD* transgene in the EyaHost system. Changing the site of tumor initiation would then request to exchange the *Eya* enhancer with another organ specific enhancer by standard transgene generation.

## Methods and Materials

### STAR★Methods

#### Key resources table

| REAGENT or RESOURCE | SOURCE | IDENTIFIER |
| --- | --- | --- |
| **Chemicals** |  |  |
| Paraformaldehyde 16% solution | Thermo Scientific Chemicals | Cat#043368.9M |
| Triton X-100 | Sigma Aldrich | Cat#T9284 |
| Vectashield mounting medium | Vector Laboratories | Cat#H-1000 |
| **Antibodies** |  |  |
| Mouse anti-pJNK (1:100) | Cell Signaling Technology | Cat#9255S; RRID:AB_2307321 |
| Rabbit anti-cleaved Dcp-1 (1:100) | Cell Signaling Technology | Cat#9578; RRID:AB_2721060 |
| Rabbit anti-pERK (1:100) | Cell Signaling Technology | Cat#4370; RRID:AB_2315112 |
| Chicken anti-beta Galactosidase (1:1000) | Abcam | Cat#ab9361; RRID:AB_307210 |
| Rat anti-elav (1:50) | DSHB | Cat#7E8A10; RRID:AB_528218 |
| Rabbit anti-scribble (1:1000) | Kind gift from Chris Q. Doe | N/A |
| Alexa Fluor 568 anti-mouse (1:500) | Thermo Fisher Scientific | Cat#A10037; RRID:AB_2534013 |
| Alexa Fluor 568 anti-Rabbit (1:500) | Thermo Fisher Scientific | Cat#A10042; RRID:AB_2534017 |
| Rhodamine anti-Chicken (1:250) | Jackson ImmunoResearch Labs | Cat#703-025-155; RRID:AB_2340348 |
| Alexa Fluor 647 anti-Rat (1:500) | Jackson ImmunoResearch Labs | Cat#712-605-153; RRID:AB_2340694 |
| Cy3 anti-guinea pig (1:500) | Jackson ImmunoResearch Labs | Cat#706-165-148; RRID:AB_2340460 |
| **Dyes** |  |  |
| Hoechst 33342 (5µg ml<sup>-1</sup>) | Thermo Fisher Scientific | Cat#H3570 |
| Alexa Fluor 488 Phalloidin (1:250) | Thermo Fisher Scientific | Cat#A12379 |
| Rhodamine Phalloidin (1:250) | Thermo Fisher Scientific | Cat#R415 |
| Alexa Fluor 647 Phalloidin (1:250) | Thermo Fisher Scientific | Cat#A22287 |
| LipidTox Red (1:250) | Thermo Fisher Scientific | Cat#H34476 |
| LipidTox Deep Red (1:250) | Thermo Fisher Scientific | Cat#H34477 |
| **Plasmids** |  |  |
| p5xQUAS-Ras<sup>V12</sup> | This study | N/A |
| p20xQUAS-shScrib | This study | N/A |
| pEya-attB-KDR | This study | N/A |
| pActin5C-KDRT>-dSTOP-KDRT>-QF2 | This study | N/A |
| p5xQUAS-mTagBFP2 | This study | N/A |
| **Experimental Models: *Drosophila melanogaster*** |  |  |
| *ey-Flp; Act-FRT-y<sup>+</sup>-FRT-Gal4, UAS-GFP; FRT82B, Tub-Gal80* | Kind gift from David Bilder |  |
| *;;FRT82B* | Bloomington Drosophila Stock Center | BDSC#5619 |
| *;UAS-Ras<sup>V12</sup>;FRT82B, scrib<sup>1</sup>* | Kind gift from Mirka Uhlirova |  |
| *;UAS-Ras<sup>V12</sup>;FRT82B, e, scrib<sup>2</sup>/TM6c* | Kind gift from Gábor Juhász |  |
| *ey-Flp;;* | Bloomington Drosophila Stock Center | BDSC#5580 |
| *;;Act-FRT-CD2-FRT-GAL4* | Bloomington Drosophila Stock Center | BDSC#4780 |
| *;;MHC-GAL4* | Bloomington Drosophila Stock Center | BDSC#55133 |
| *;;Mef2-GAL4* | Bloomington Drosophila Stock Center | BDSC#27390 |
| *;;R4-GAL4* | Bloomington Drosophila Stock Center | BDSC#33832 |
| *;;da-GAL4* | Bloomington Drosophila Stock Center | BDSC#55850 |
| *;;He-GAL4* | Bloomington Drosophila Stock Center | BDSC#8699 |
| *Tub-Gal80<sup>ts</sup>* | Bloomington Drosophila Stock Center | BDSC#7018; BDSC#7108 |
| *;;UAS-mCherry<sup>RNAi</sup>* | Bloomington Drosophila Stock Center | BDSC#35785 |
| *;UAS-RFP<sup>RNAi</sup>* | Bloomington Drosophila Stock Center | BDSC#67852 |
| *;;UAS-Luc<sup>RNAi</sup>* | Bloomington Drosophila Stock Center | BDSC#35788 |
| *;;UAS-white<sup>RNAi</sup>* | Bloomington Drosophila Stock Center | BDSC#33762 |
| *w<sup>[1118]</sup> GD ctrl* | Vienna Drosophila Research Center | VDRC#60000 |
| *;UAS-atg1<sup>RNAi</sup>* | Vienna Drosophila Research Center | VDRC#16133 |
| *;;UAS-pten<sup>RNAi</sup>* | Bloomington Drosophila Stock Center | BDSC#33643 |
| *;;kibra-LacZ* | Bloomington Drosophila Stock Center | BDSC#11567 |
| *;;10xSTAT92E-GFP* | Bloomington Drosophila Stock Center | BDSC#26197 |
| *;atg8-atg8::3xCherry* | Kind gift from Gábor Juhász | N/A |
| *UAS-CD8::Cherry* | Bloomington Drosophila Stock Center | BDSC#27391; BDSC#27392 |
| *QUAS-mCD8::GFP;;* | Bloomington Drosophila Stock Center | BDSC#30001 |
| *;;20xQUAS-Gal80* | This study | N/A |
| *;;5xQUAS-Gal80* | Bloomington Drosophila Stock Center | BDSC#51950 |
| *;;QUAS-Gal4* | Bloomington Drosophila Stock Center | BDSC#83132 |
| *5xQUAS-Ras<sup>V12</sup>;;* | This study | N/A |
| *;20xQUAS-scrib<sup>RNAi</sup>* | This study | N/A |
| *Act-KDRT-STOP-KDRT-QF2;;* | This study | N/A |
| *;Eya-KDR* | This study | N/A |
| *5xQUAS-mTagBFP2;;* | This study | N/A |
| *Act-KDRT-STOP-KDRT-QF2, QUAS-mCD8::GFP, ey-flp;;* | This study | N/A |
| *5xQUAS-Ras<sup>V12</sup>; 20xQUAS-scrib<sup>RNAi</sup>; Dr/TM6c* | This study | N/A |
| *5xQUAS-Ras<sup>V12</sup>; 20xQUAS-scrib<sup>RNAi</sup>; Act-FRT-CD2-FRT-GAL4, QUAS-Gal80/TM6c* | This study | N/A |
| *5xQUAS-Ras<sup>V12</sup>; 20xQUAS-scrib<sup>RNAi</sup>; da-gal4* | This study | N/A |
| *5xQUAS-Ras<sup>V12</sup>; 20xQUAS-scrib<sup>RNAi</sup>; MHC-Gal4, Tub-Gal80<sup>ts</sup>* | This study | N/A |
| *5xQUAS-Ras<sup>V12</sup>; 20xQUAS-scrib<sup>RNAi</sup>; Mef2-GAL4* | This study | N/A |
| *5xQUAS-Ras<sup>V12</sup>; 20xQUAS-scrib<sup>RNAi</sup>; R4-Gal4, Tub-Gal80<sup>ts</sup>* | This study | N/A |
| *5xQUAS-Ras<sup>V12</sup>; 20xQUAS-scrib<sup>RNAi</sup>; He-Gal4* | This study | N/A |
| *5xQUAS-Ras<sup>V12</sup>; 20xQUAS-scrib<sup>RNAi</sup>; QUAS-Gal4* | This study | N/A |
| **Standard potato mash fly food recipe (per litre)** |  |  |
| 32.7 g dried potato powder | Mills AS | N/A |
| 60 g sucrose | Dansukker | N/A |
| 27.3 g dry yeast | Pals AS | Cat#20610 |
| 7.3 g agar | Pals AS | Cat#77000 |
| 4.55 ml propionic acid | Sigma Aldrich | Cat#402907 |
| 2 g nipagin in 50ml absolute alcohol | Sigma Aldrich | Cat#H5501 |
| **Imaging** |  |  |
| Leica Thunder Imager Model Organism | Leica Microsystems | N/A |
| Zeiss LSM880 Confocal microscope | Zeiss | N/A |
| **Software** |  |  |
| Fiji | https://fiji.sc/ | N/A |
| Adobe Illustrator | Adobe | N/A |
| Adobe Photoshop | Adobe | N/A |
| ZEN Microscopy Software | Zeiss | N/A |
| Prism 10 | GraphPad | N/A |

#### Resource availability

For resource requests, contact.

#### Method details

##### Plasmid constructs for transgenesis

Plasmids used for generating transgenic flies are listed in the STAR★Methods. The listed constructs were made by conventional restriction enzyme-based cloning and Gibson Assembly (New England Biolabs). Oligonucleotides for PCR were purchased from Eurofins or MicroSynth. Plasmid constructs were first verified by restriction enzyme digestion, followed by DNA sequencing using Eurofins or MicroSynth services.

Briefly, pQUAS-WALIUM20 (https://dgrc.bio.indiana.edu//stock/1474) was used as the p5xQUAS backbone, while p20xQUAS backbone was generated by subcloning 15xQUAS enhancer from pQUASp (https://www.addgene.org/46162/) into p5xQUAS-WALIUM20 with SphI-site using PCR amplification. shScrib2 and mTagBFP2 fragments were further subcloned into respective pQUAS backbones to generate p20xQUAS-shScrib2 and p5xQUAS-mTagBFP2 respectively using PCR amplification, while p5xQUAS-RasV12 was a kind gift from Peter Gallant.

pActin5C-KDRT>-dSTOP-KDRT>-QF2 was generated by first making pActin5C-KDRT-dSTOP-KDTR-myrRFP from pJFRC164-21XUAS-KDRT>-dSTOP-KDRT>-myr::RFP (https://www.addgene.org/32141/) by replacing 21xUAS with pActin5C, followed by amplify QF2 from pQF2wWB (https://www.addgene.org/61313/) and subcloning it into pActin5C-KDRT-dSTOP-KDTR-myrRFP vector by replacing myrRFP with QF2.

##### Transgenesis

PhiC31 integrase-mediated site-specific transgenesis and genomic PCR confirmation of insertion was performed at BestGene (https://www.thebestgene.com). Successful recombinants were screened for mini-white^+^ in the white mutant background. *Act-KDRT-STOP-KDRT-QF2* and *5xQUAS-Ras^V12^* was inserted in Attp18 (BDSC#32107), *Eya-KDR* in Attp14, *20xQUAS-scribRNAi* in attp5su(Hw) (BDSC#34765) and attp40, and *5xQUAS- mTagBFP2* in Attp4, 20xQUAS-Gal80 in VK00027 (BDSC#9744).

##### Fly husbandry and experimental crosses

Flies were cultured in standard potato mash fly food with 24hr egg lays to stage the animals and under controlled humidity and temperature (25°C). For Tub-Gal80ts experiments, eggs were laid and developed at 18°C for 6-8 days and transferred to 29°C to induce RNAi (R4 tester); eggs were collected for 24 hours at 25°C and scored at 10 days after transferred to 29 C (MHC tester). Eggs were collected and developed for 8-10 days at 18 degrees and analyzed at 11 days after being transferred to 29 degrees (Mef2 tester). To obtain lingering animals, vial crowdedness levels has to be controlled. We observed that a higher density of larvae promoted the delay of pupation and lingering past day 5 AEL. For this, we optimized virgin collections to 5 days with yeast-supplemented food to boost egg-laying before mating. Crosses were then set with 8 virgins and 6 males to achieve an optimal egg-lay and vial crowdedness. For the pupariation assay, only larva and pupa with the correct phenotype were counted from day 6 until 10 after egg-lay.

##### Drosophila stocks and genetics

For a complete overview of all fly stocks of the EyaHOST system, please check suppl. table 1. Additionally, for a list of all genotypes per figure, please refer to suppl. table 2. The following fly strains have been obtained from Bloomington Drosophila Stock Center *;;FRT82B* (BDSC#5619), *ey-Flp;;* (BDSC#5580), *;;Act-FRT-CD2-FRT-GAL4* (BDSC#4780), *;;MHC-GAL4* (BDSC#55133), *;;Mef2-GAL4* (BDSC#27390), *;;R4-GAL4* (BDSC#33832*), ;;da-GAL4* (BDSC#55850), *;;He-GAL4* (BDSC#8699), *Tub-Gal80^ts^* (BDSC#7018; BDSC#7108), *;;UAS-atg1^RNAi^* (BDSC#26731), *;;UAS-mCherry^RNAi^* (BDSC#35785), *;UAS-RFP^RNAi^* (BDSC#67852), *;;UAS-Luc^RNAi^* (BDSC#35788), ;;UAS-white^RNAi^ (BDSC#33762), *;;kibra-LacZ* (BDSC#11567), *;;10xSTAT92E-GFP* (BDSC#26197), *UAS-CD8::Cherry* (BDSC#27391; BDSC#27392), *QUAS-mCD8::GFP;;* (BDSC#30001), *;;QUAS-Gal80* (BDSC#51950), *;;QUAS-Gal4* (BDSC#83132). From the Vienna Drosophila stock center we used lines *w^[1118]^* GD ctrl (VDRC#60000), *;UAS-atg1^RNAi^* (VDRC#16133). The following stocks were kindly gifted *ey-Flp; Act-FRT-y+- FRT-Gal4, UAS-GFP; FRT82B, Tub-Gal80* (D. Bilder), *;UAS-RasV12;FRT82B, scrib^1^* (M. Uhlirova), *;UAS-RasV12;FRT82B, e, scrib^2^/TM6c, ;3xCherry::Atg8a* (G. Juhász).

##### Sample preparation and histology

*Drosophila* larva with the correct genotype were selected and washed in PBS. For EAD/tumor preparations, partial dissection was first performed by cutting the larva in half in chilled PBS, inverting the carcass, and removing the adjacent tissue. Samples were then fixed in 4% PFA (Paraformaldehyde dissolved in 1XPBS) in 1.5ml tubes for 30 minutes and washed in 0.5% PBT at RT. Subsequently, antibody or dye staining was performed overnight at 4C and washed in 0.05% PBT (Triton X-100 dissolved in 1xPBS). Samples were then completely dissected and mounted in a glass slide with a spacer and Vectashield mounting medium. Antibody/Dye list and concentrations used are available in the STAR★Methods table. Fat bodies were similarly dissected, but instead washed inside plastic cages with nylon mesh to avoid tissue disruption. Muscle fillets were dissected in silicon plates by pinning the larva and cutting the larva dorsally through the midline. Adjacent tissue was then removed, and the body was pinned laterally, followed by the steps described above. Samples were imaged with a Zeiss LSM 880 confocal using a 10x or 20x air objective. For volumetric analysis, z-stacks were acquired from the top to the bottom of the sample with a 3 or 4 µm z-slice interval. For whole animal imaging, larvae were washed in PBS and then heat-shocked at 75C in glycerol for euthanization, followed by immediate imaging with Leica Thunder Imager Model Organism.

##### Quantification and statistical analysis

For volumetric analysis of tumor size, we performed 3D reconstruction of tumor confocal z-stacks by semi-automatic thresholding of GFP signal in Fiji. Images were subjected to Gaussian blur for denoising and to speed-up analysis. The best-fitting thresholding method was qualitatively assessed through visualization of the true signal and mask overlay. GraphPad was used for statistical analysis. The data was first checked for normality using the Shapiro-Wilk test. Statistical tests used to assess for statistical differences between groups are indicated in the figure legend. Error bars stand for SD, unless noted otherwise. A minimum of two repeats were done for all experiments.

#### Data and Code Availability

Raw data is available upon request.

## Acknowledgements

We thank R. Sousa Nunes and Justin Kumar for discussions on the construction of genetic tools for EyaHOST. We thank C. Potter, J.Kumar, P. Gallant, Bloomington Stock Centre, the TRiP at Harvard Medical School (NIH/NIGMS R01-GM084947), VDRC, and the Developmental Studies Hybridoma Bank for fly stocks and reagents This work was supported in part by The Norwegian Research Council Toppforsk and Center of Excellence, CanCell grants #262652 and #276070. We also thank the Core Facilities for Advanced Light Microscopy and Advanced Electron Microscopy at Oslo University Hospital.

## Author contributions

Experimental design C.D., J.T.R, A.J. and T.E.R.,; Methodology C.D., J.T.R, A.J. D.L., R.K., S.M.; Investigation C.D, J.T.R, A.J., D.L., R.K., S.M., A.A.G.; Data analysis C.D, J.T.R.,A.J.; Manuscript writing C.D, J.T.R, T.E.R.; Supervision C.D. and T.E.R; Materials T.E.R; Funding T.E.R.

## Declaration of interests

The authors have no conflict of interest.

Supplemental Information

Supplemental table 1 – Overview of all the fly lines of the EyaHOST system.

Supplemental table 2 – Description of the full genotypes per figure.

---

**Suppl. Fig. 1.**
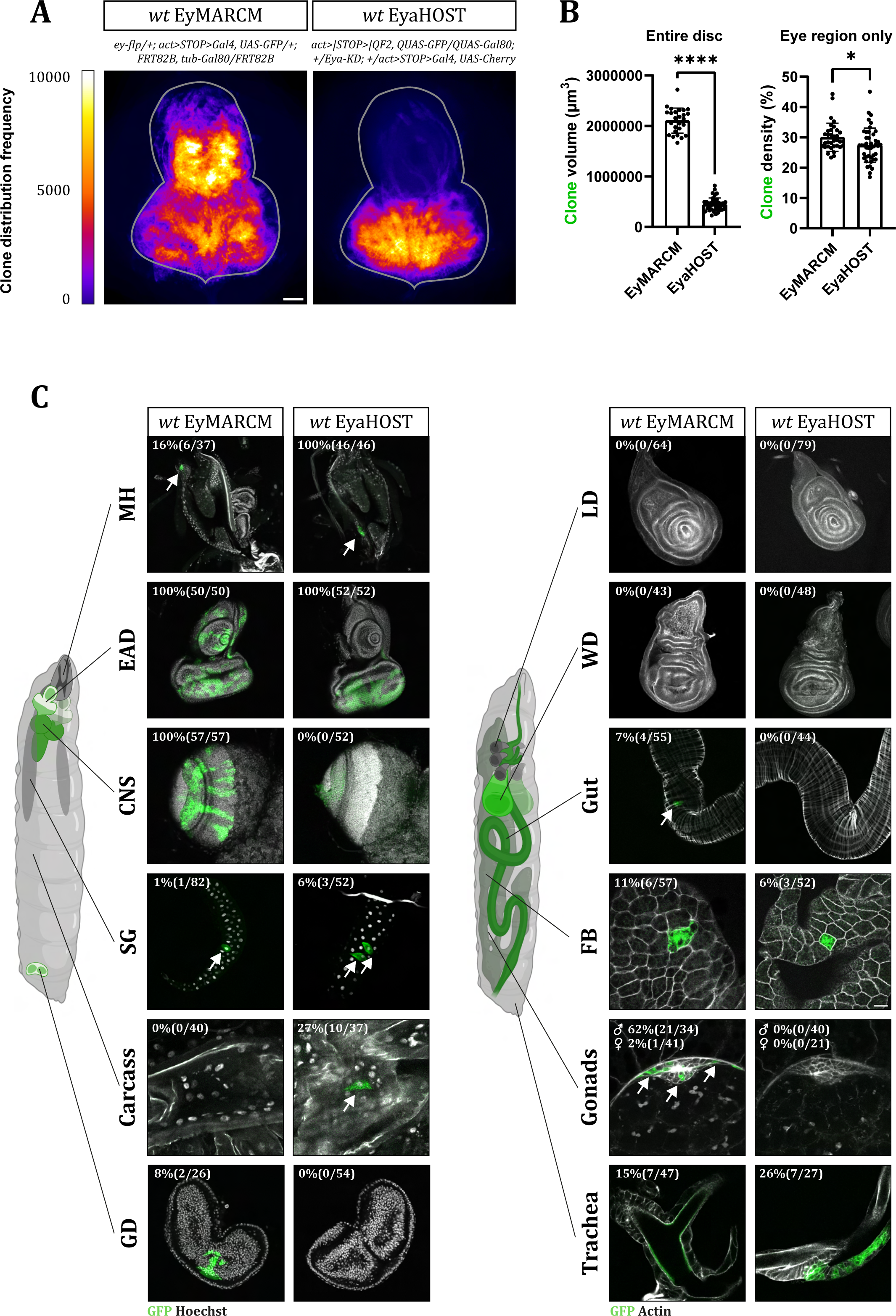
Characterization of the clonal patterns of the EyaHOST system. (A) WT clone frequency heatmap, generated by superimposed EADs, shows the spatial distribution of activation within the EyMARCM and EyaHOST systems. Scale bar - 50 μm (B) Quantification of total EAD WT clone volume (left) and WT clone density in the eye region only (right) for both EyMARCM and EyaHOST systems. Statistical significance between groups was determined using the Mann-Whitney U test: *p ≤ 0.0332, ****p ≤ 0.000. (C) Multi-organ characterization of EyMARCM and EyaHOST activation frequencies at the 3rd instar larval stage. The organs analyzed include: (left) Mouth hook (MH), eye-antennal disc (EAD), central nervous system (CNS), salivary gland (SG), carcass, and genital discs (GD); and (right) leg discs (LD), wing discs (WD), gut, fat body (FB), gonads, and trachea. Organs were counterstained with Hoechst for nuclei or phalloidin for actin (gray). Quantifications shown for the frequency of observation of GFP- positive cells (green) in each organ. Organs were counterstained with Hoechst for nuclei or phalloidin for actin (gray).

**Suppl. Fig. 2.**
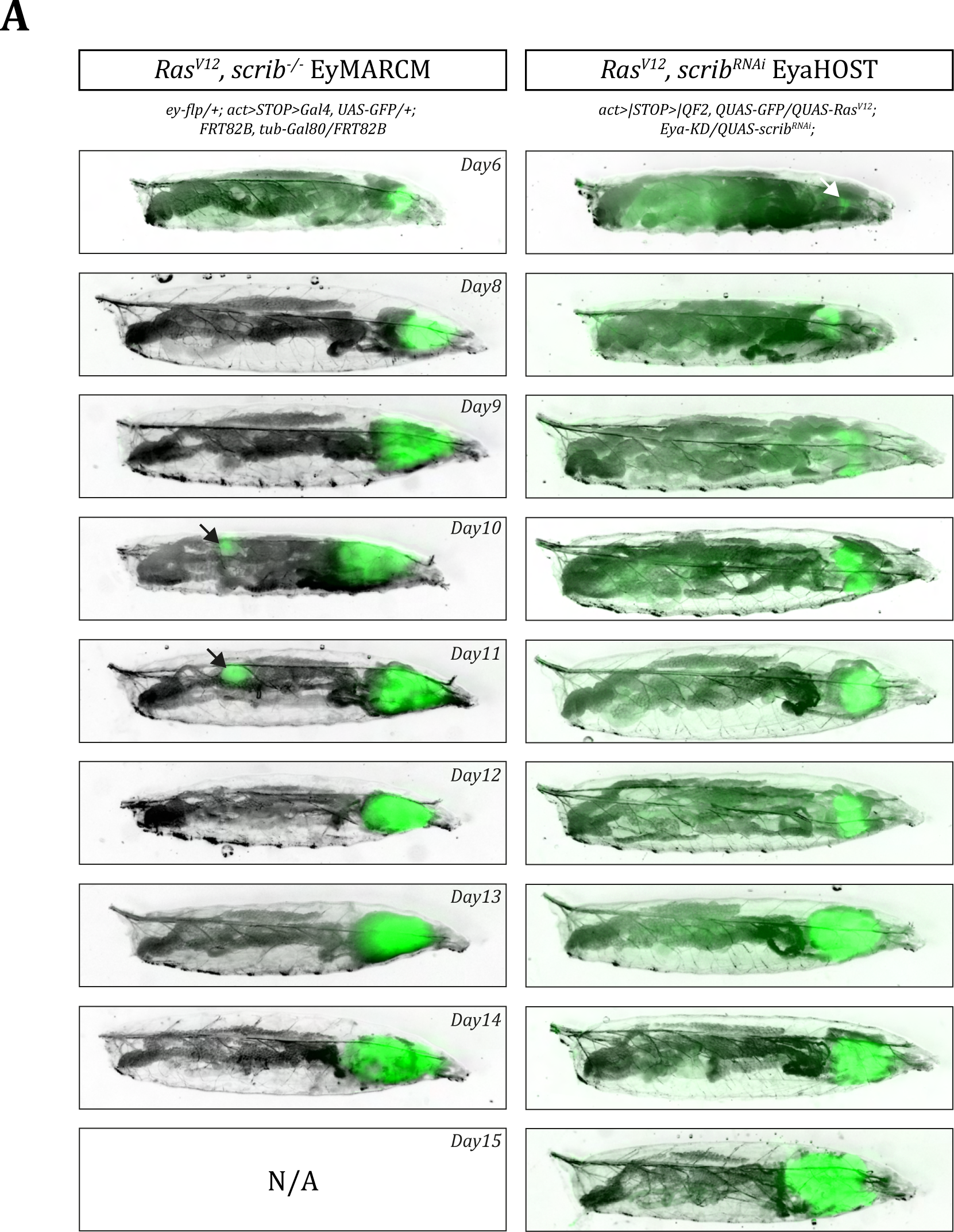
Temporal evolution of Ras^V12^, scrib tumour volume progression in the EyMARCM and EyaHOST systems. (A) Whole-animal stereoscope images tracking the size of GFP-positive Ras^V12^, scrib tumours (green) from D6 to D15. No image is shown for D15 EyMARCM due to no survival. Black arrows indicate commonly observed gonad tumours in EyMARCM system.

**Suppl. Fig. 3.**
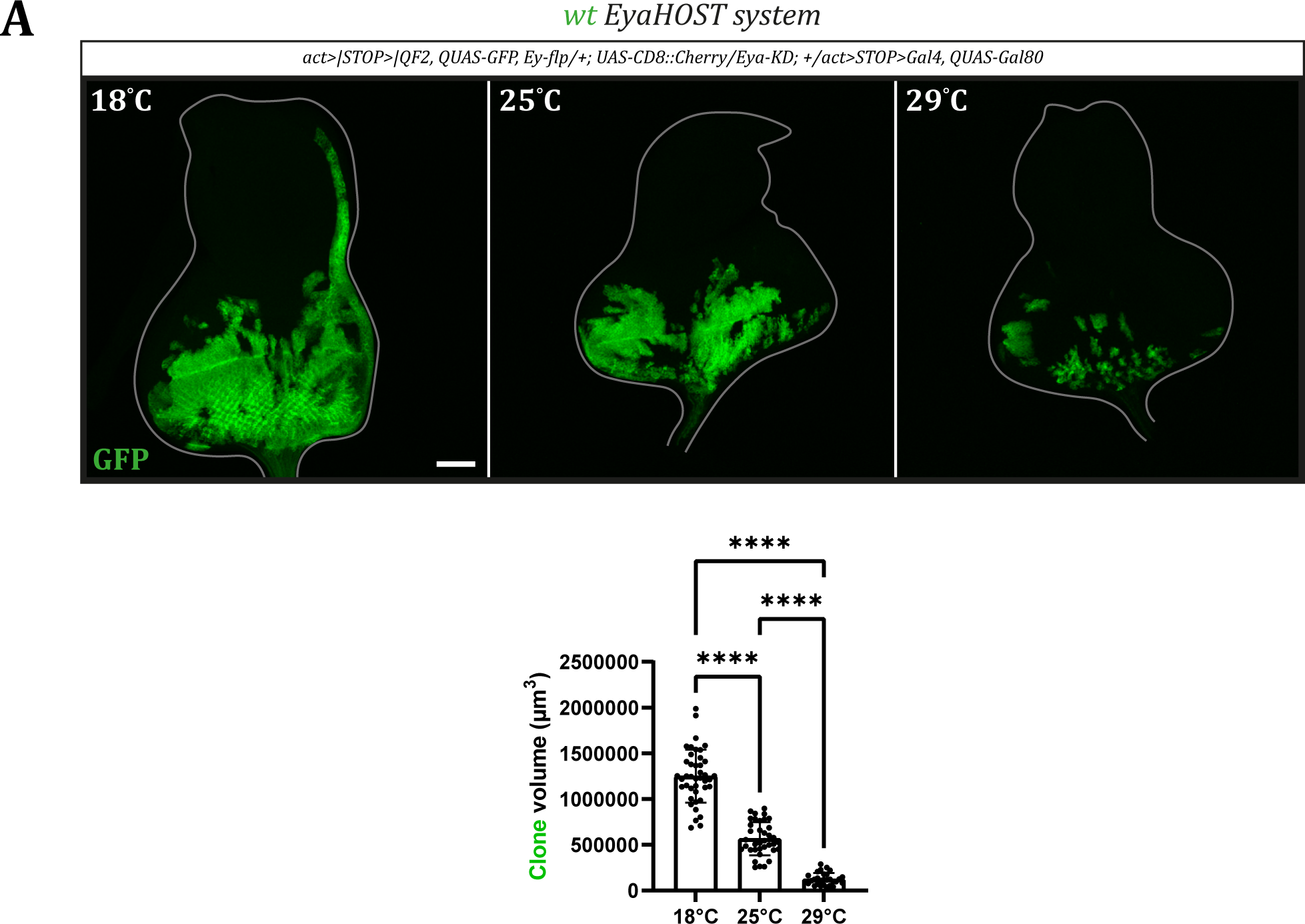
The EyaHOST system enables targeted gene knockdown specifically in neighbouring surrounding cells. (A) EyaHOST system adult *Drosophila* eye displaying the activation domains of the QF-QUAS (GFP positive – green) and GAL4-UAS (Cherry positive – red) compartments. The specific knockdown of the retina pigmentation gene *white* in neighbouring cells is indicated by the absence of pigmentation (light).

**Suppl. Fig. 4.**
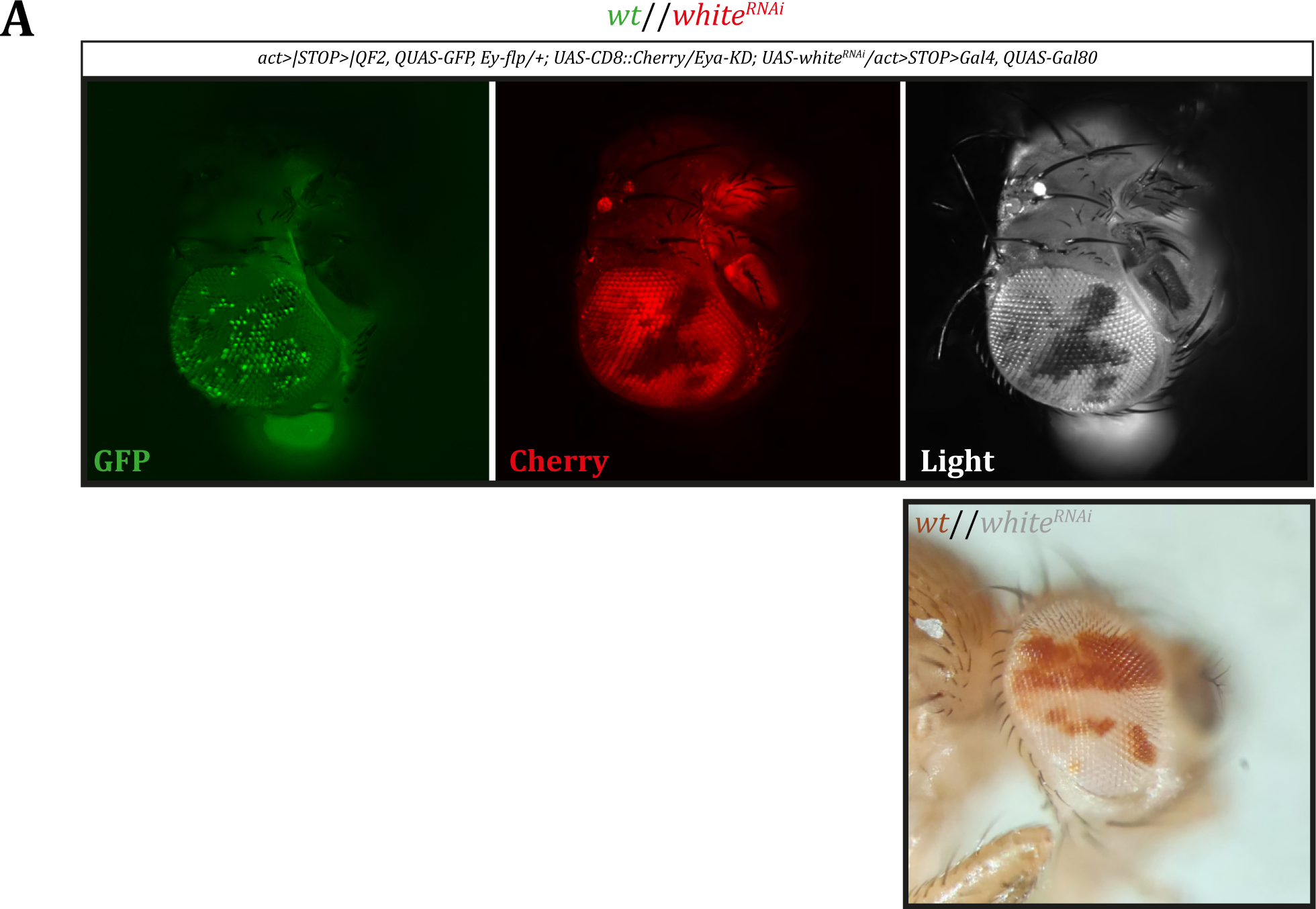
The activation of the EyaHOST system in the EAD is temperature sensitive. (A) Confocal images of EAD displaying the clonal activation of the EyaHOST system at different temperatures (18°C, 25°C, and 29°C). Quantification of clone volume at different temperatures are shown. Statistical differences between groups were calculated using a Kruskal-Wallis and post-hoc Dunn’s multiple comparison test: ****p ≤ 0.000. Scale bar - 50 μm.

**Suppl. Fig. 5.**
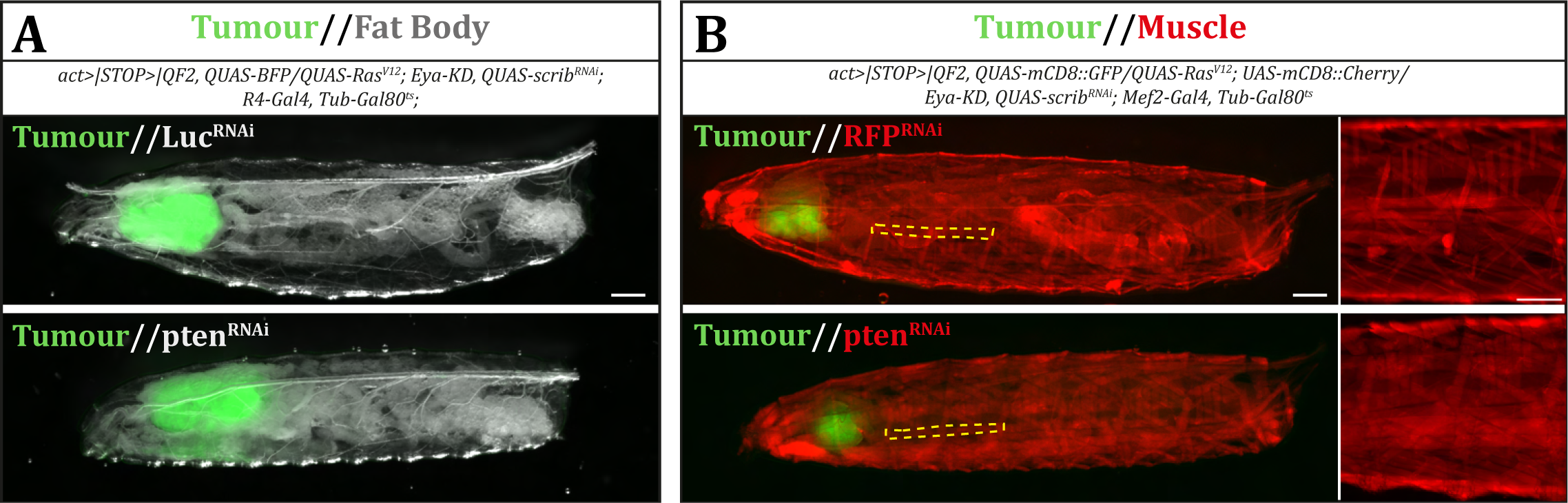
PTEN knockdown in peripheral organs rescues tissue atrophy. Whole animal stereoscope image are shown for control animals and for pten knockdown in fat body (A) or muscle (B). Tumours are labelled in green, fat body is observable through light, and muscles where labelled with Cherry (red). Scale bar – 700 μm.

