## Supplementary table 2 for "EyaHOST, a modular genetic system for investigation of intercellular and tumor-host interactions *in Drosophila melanogaster*"

<table>
<tr><th>Figure</th><th>Genotype</th></tr>
<tr><td>1B</td><td>ey-flp/+;Act>STOP>Gal4, UAS-GFP/+;FRT82B, tub-Gal80/FRT82B</td></tr>
<tr><td>1C</td><td>Act>|STOP>|QF2, QUAS-CD8::GFP, ey-flp/QUAS-Gal80; +/eya-KDR; +/Act>STOP>Gal4, UAS-CD8::Cherry</td></tr>
<tr><td>1D</td><td>Act>|STOP>|QF2, QUAS-CD8::GFP/QUAS-Ras$^{V12}$; eya-KDR/QUAS-scrib$^{RNAi}$;</td></tr>
<tr><td>1E</td><td>Act>|STOP>|QF2, QUAS-CD8::GFP/QUAS-Ras$^{V12}$; eya-KDR/QUAS-scrib$^{RNAi}$;</td></tr>
<tr><td rowspan="3">2A</td><td>ey-flp/+; Act>STOP>Gal4, UAS-GFP/UAS-Ras$^{V12}$; FRT82B, tub-Gal80/FRT82B, scrib$^{1}$</td></tr>
<tr><td>ey-flp/+; Act>STOP>Gal4, UAS-GFP/UAS-Ras$^{V12}$; FRT82B, tub-Gal80/FRT82B, e, scrib$^{2}$</td></tr>
<tr><td>Act>|STOP>|QF2, QUAS-CD8::GFP/QUAS-Ras$^{V12}$; eya-KDR/QUAS-scrib$^{RNAi}$;</td></tr>
<tr><td rowspan="3">2B</td><td>ey-flp/+; Act>STOP>Gal4, UAS-GFP/UAS-Ras$^{V12}$; FRT82B, tub-Gal80/FRT82B, scrib$^{1}$</td></tr>
<tr><td>ey-flp/+; Act>STOP>Gal4, UAS-GFP/UAS-Ras$^{V12}$; FRT82B, tub-Gal80/FRT82B, e, scrib$^{2}$</td></tr>
<tr><td>Act>|STOP>|QF2, QUAS-CD8::GFP/QUAS-Ras$^{V12}$; eya-KDR/QUAS-scrib$^{RNAi}$;</td></tr>
<tr><td rowspan="2">2C</td><td>Act>|STOP>|QF2, QUAS-CD8::GFP/+; eya-KDR/+;</td></tr>
<tr><td>Act>|STOP>|QF2, QUAS-CD8::GFP/QUAS-Ras$^{V12}$; eya-KDR/QUAS-scrib$^{RNAi}$;</td></tr>
<tr><td rowspan="2">2D</td><td>Act>|STOP>|QF2, QUAS-CD8::GFP/+; eya-KDR/+;</td></tr>
<tr><td>Act>|STOP>|QF2, QUAS-CD8::GFP/QUAS-Ras$^{V12}$; eya-KDR/QUAS-scrib$^{RNAi}$;</td></tr>
<tr><td rowspan="2">2E</td><td>Act>|STOP>|QF2, QUAS-CD8::GFP/+; eya-KDR/+;</td></tr>
<tr><td>Act>|STOP>|QF2, QUAS-CD8::GFP/QUAS-Ras$^{V12}$; eya-KDR/QUAS-scrib$^{RNAi}$;</td></tr>
<tr><td rowspan="2">3A</td><td>Act>|STOP>|QF2, QUAS-CD8::GFP/+; eya-KDR/+;</td></tr>
<tr><td>Act>|STOP>|QF2, QUAS-CD8::GFP/QUAS-Ras$^{V12}$; eya-KDR/QUAS-scrib$^{RNAi}$;</td></tr>
<tr><td rowspan="2">3B</td><td>Act>|STOP>|QF2, QUAS-mTagBFP2, ey-flp/+; 10xSTAT-GFP/eya-KDR; +/Act>STOP>Gal4, QUAS-Gal80</td></tr>
<tr><td>Act>|STOP>|QF2, QUAS-mTagBFP2, ey-flp/QUAS-Ras$^{V12}$; 10xSTAT-GFP/QUAS-scrib$^{RNAi}$, eya-KDR; +/Act>STOP>Gal4, QUAS-Gal80</td></tr>
<tr><td rowspan="2">3C</td><td>Act>|STOP>|QF2, QUAS-CD8::GFP, ey-flp/+; +/eya-KDR; kibra-LacZ/Act>STOP>Gal4, QUAS-Gal80</td></tr>
<tr><td>Act>|STOP>|QF2, QUAS-CD8::GFP, ey-flp/QUAS-Ras$^{V12}$; +/ QUAS-scrib$^{RNAi}$, eya-KDR; kibra-LacZ/Act>STOP>Gal4, QUAS-Gal80</td></tr>
<tr><td rowspan="2">4A</td><td>Act>|STOP>|QF2, QUAS-CD8::GFP, ey-flp/+; UAS-CD8::Cherry/eya-KDR; +/Act>STOP>Gal4, QUAS-Gal80</td></tr>
<tr><td>Act>|STOP>|QF2, QUAS-CD8::GFP, ey-flp/QUAS-Ras$^{V12}$; UAS-CD8::Cherry/QUAS-scrib$^{RNAi}$, eya-KDR; +/Act>STOP>Gal4, QUAS-Gal80</td></tr>
<tr><td rowspan="2">4B</td><td>Act>|STOP>|QF2, QUAS-CD8::GFP, ey-flp/+; +/eya-KDR; +/He-Gal4,UAS-CD8::Cherry</td></tr>
<tr><td>Act>|STOP>|QF2, QUAS-CD8::GFP, ey-flp/QUAS-Ras$^{V12}$; +/QUAS-scrib$^{RNAi}$, eya-KDR; +/He-Gal4,UAS-CD8::Cherry</td></tr>
<tr><td>4C</td><td>Act>|STOP>|QF2, QUAS-CD8::GFP/QUAS-Ras$^{V12}$; eya-KDR/QUAS-scrib$^{RNAi}$; UAS-CD8::Cherry/MHC-Gal4, tub-Gal80$^{ts}$</td></tr>
<tr><td>4D</td><td>Act>|STOP>|QF2, QUAS-CD8::GFP/QUAS-Ras$^{V12}$; eya-KDR/QUAS-scrib$^{RNAi}$; UAS-CD8::Cherry/R4-Gal4, tub-Gal80$^{ts}$</td></tr>
<tr><td>4E</td><td>Act>|STOP>|QF2, QUAS-CD8::GFP, ey-flp/QUAS-Ras$^{V12}$; UAS-CD8::Cherry/QUAS-scrib$^{RNAi}$, eya-KDR; +/da-Gal4</td></tr>
<tr><td>4F</td><td>Act>|STOP>|QF2, QUAS-CD8::GFP, ey-flp/QUAS-Ras$^{V12}$; UAS-CD8::Cherry/QUAS-scrib$^{RNAi}$, eya-KDR; +/QUAS-Gal4</td></tr>
<tr><td>5A</td><td>Act>|STOP>|QF2, QUAS-CD8::GFP, ey-flp/+; Atg8-Atg8::3xCherry/eya-KDR; +/Act>STOP>Gal4, QUAS-Gal80</td></tr>
</table>

<table>
<tr><td></td><td>Act>|STOP>|QF2, QUAS-CD8::GFP, ey-flp/QUAS-Ras$^{V12}$; Atg8-Atg8::3xCherry/ QUAS-scrib$^{RNAi}$, eya-KDR; +/Act>STOP>Gal4, QUAS-Gal80</td></tr>
<tr><td rowspan="2">5B</td><td>Act>|STOP>|QF2, QUAS-CD8::GFP/QUAS-Ras$^{V12}$; +/QUAS-scrib$^{RNAi}$, eya-KDR; UAS-mCD8::Cherry /R4-Gal4, tub-Gal80$^{ts}$</td></tr>
<tr><td>Act>|STOP>|QF2, QUAS-CD8::GFP/QUAS-Ras$^{V12}$; UAS-Atg1$^{RNAi}$/QUAS-scrib$^{RNAi}$, eya-KDR; UAS-mCD8::Cherry/R4-Gal4, tub-Gal80$^{ts}$</td></tr>
<tr><td rowspan="2">5C</td><td>Act>|STOP>|QF2, QUAS-CD8::GFP/QUAS-Ras$^{V12}$; +/QUAS-scrib$^{RNAi}$, eya-KDR; UAS-mCD8::Cherry /MHC-Gal4, tub-Gal80$^{ts}$</td></tr>
<tr><td>Act>|STOP>|QF2, QUAS-CD8::GFP/QUAS-Ras$^{V12}$; UAS-Atg1$^{RNAi}$/QUAS-scrib$^{RNAi}$, eya-KDR; UAS-mCD8::Cherry/MHC-Gal4, tub-Gal80$^{ts}$</td></tr>
<tr><td rowspan="2">6A</td><td>Act>|STOP>|QF2, QUAS-CD8::GFP, ey-flp/+;+/eya-KDR; +/Act>STOP>Gal4, QUAS-Gal80</td></tr>
<tr><td>Act>|STOP>|QF2, QUAS-CD8::GFP, ey-flp/ QUAS-Ras$^{V12}$;UAS-p35/QUAS-scrib$^{RNAi}$, eya-KDR; +/Act>STOP>Gal4, QUAS-Gal80</td></tr>
<tr><td rowspan="2">Suppl. 1A,B</td><td>ey-flp/+;Act>STOP>Gal4, UAS-GFP/+;FRT82B, tub-Gal80/FRT82B</td></tr>
<tr><td>Act>|STOP>|QF2, QUAS-CD8::GFP, ey-flp/QUAS-Gal80; +/eya-KDR; +/ Act>STOP>Gal4, UAS-CD8::Cherry</td></tr>
<tr><td rowspan="2">Suppl. 1C</td><td>ey-flp/+;Act>STOP>Gal4, UAS-GFP/+;FRT82B, tub-Gal80/FRT82B</td></tr>
<tr><td>Act>|STOP>|QF2, QUAS-CD8::GFP/+; eya-KDR/+;</td></tr>
<tr><td rowspan="2">Suppl. 2A</td><td>ey-flp/+; Act>STOP>Gal4, UAS-GFP/UAS-Ras$^{V12}$; FRT82B, tub-Gal80/FRT82B, scrib$^{1}$</td></tr>
<tr><td>Act>|STOP>|QF2, QUAS-CD8::GFP/QUAS-Ras$^{V12}$; eya-KDR/QUAS-scrib$^{RNAi}$;</td></tr>
<tr><td>Suppl. 3A</td><td>Act>|STOP>|QF2, QUAS-CD8::GFP, ey-flp/+;UAS-CD8::Cherry/eya-KDR; +/Act>STOP>Gal4, QUAS-Gal80</td></tr>
<tr><td>Suppl. 4A</td><td>Act>|STOP>|QF2, QUAS-CD8::GFP, ey-flp/+;UAS-CD8::Cherry/eya-KDR;UAS-white$^{RNAi}$/Act>STOP>Gal4, QUAS-Gal80</td></tr>
<tr><td rowspan="2">Suppl. 5A</td><td>Act>|STOP>|QF2, QUAS-mTagBFP2/QUAS-Ras$^{V12}$; +/QUAS-scrib$^{RNAi}$, eya-KDR; UAS-Luc$^{RNAi}$/R4-Gal4, tub-Gal80$^{ts}$</td></tr>
<tr><td>Act>|STOP>|QF2, QUAS-mTagBFP2/QUAS-Ras$^{V12}$; +/QUAS-scrib$^{RNAi}$, eya-KDR; UAS-pten$^{RNAi}$/R4-Gal4, tub-Gal80$^{ts}$</td></tr>
<tr><td rowspan="2">Suppl. 5B</td><td>Act>|STOP>|QF2, QUAS-CD8::GFP/QUAS-Ras$^{V12}$; UAS-RFP$^{RNAi}$/QUAS-scrib$^{RNAi}$, eya-KDR; UAS-mCD8::Cherry/Mef2-Gal4</td></tr>
<tr><td>Act>|STOP>|QF2, QUAS-CD8::GFP/QUAS-Ras$^{V12}$; UAS-mCD8::Cherry/QUAS-scrib$^{RNAi}$, eya-KDR; UAS-pten$^{RNAi}$/Mef2-Gal4</td></tr>
</table>
